# A Role for Astrocyte Metabolism in Species-Specific Neuronal Development

**DOI:** 10.64898/2026.07.15.737608

**Authors:** Sheila Steiner, Kaia Foster, Rebecca Chinn, Joshua Pratt, Sarah Fernandes, Amandeep Sharma, Renata Santos, Christian Metallo, Maria Carolina Marchetto, Fred H. Gage

## Abstract

Human neurons develop more slowly than non-human primate (NHP) neurons, a phenomenon called neoteny, but research has primarily focused on neuron-intrinsic drivers. We hoped to further elucidate any species-specific divergence in function and the astrocytes’ role in influencing species-specific neurodevelopment rate. In this study, we identified a delayed onset of gliogenesis in human versus NHP organoid models. Transcriptomic and ^13^C metabolic flux analyses of iPSC-derived astrocytes revealed distinct metabolic profiles: NHP astrocytes exhibit increased serine and glycine synthesis, whereas human astrocytes show elevated lactate secretion, suggesting a change in the metabolic role of astrocytes across primate evolution. We then assessed the impact of these different species’ astrocytes on neuronal development. We observed an increase in electrophysiological maturation and a change in transcriptomic neuronal development trajectory in human neurons cultured with rhesus astrocyte conditioned media as opposed to human astrocyte conditioned media. Human astrocyte secretomes were enriched for synaptogenic and axon-growth proteins, which could indicate they play a greater role in supporting structural complexity and dendritic arborization over rapid maturation when compared to NHP astrocytes. Finally, chemical inhibition of PHGDH demonstrated that these changes in neuronal differentiation are partially mediated by the different metabolic roles that astrocytes play in humans versus NHPs. Collectively, our results reveal a cell-extrinsic role for astrocyte metabolism in shaping human-specific neurodevelopmental timing and trajectories.

**Graphical Abstract:** 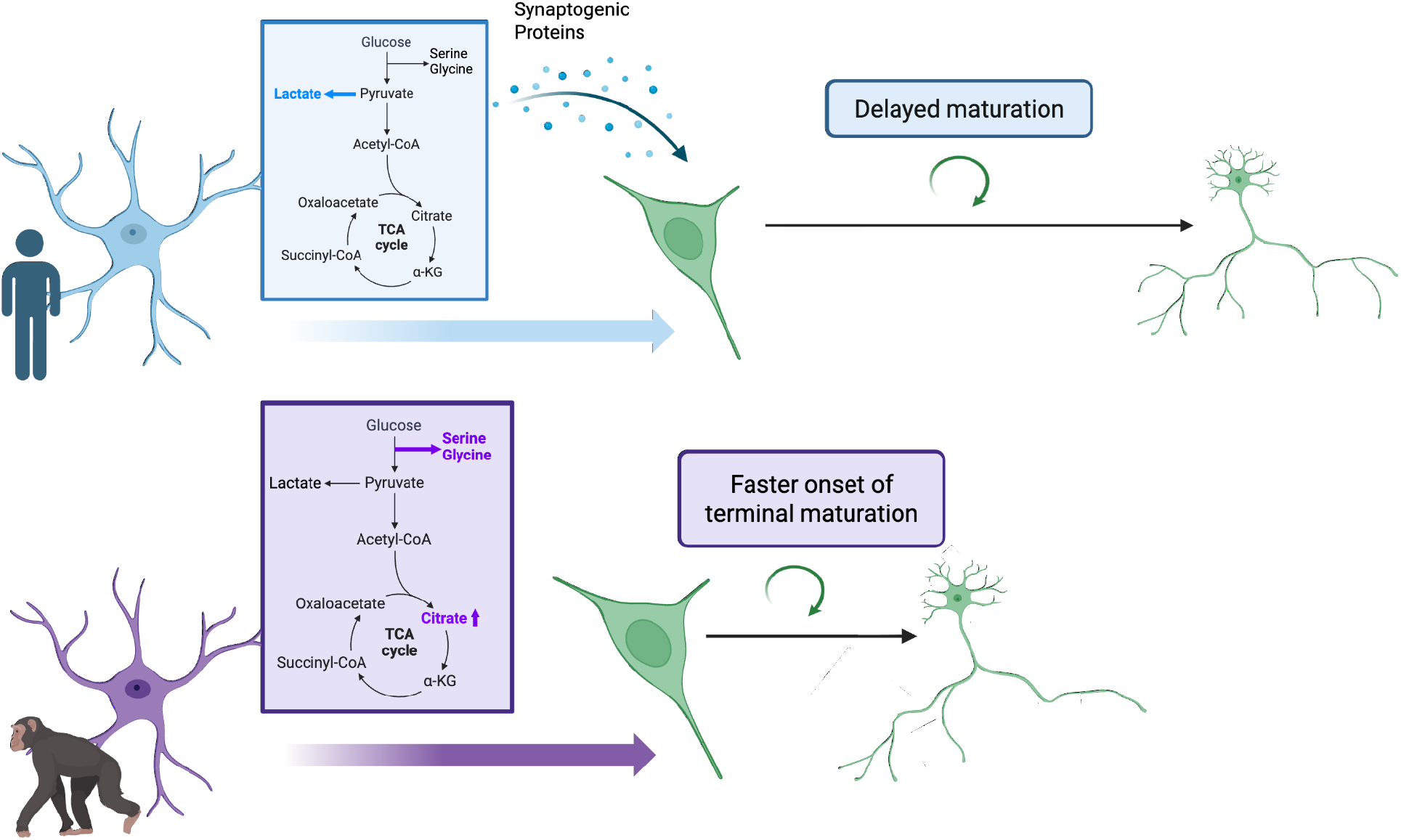

## Introduction

Astrocytes are fundamental to the function and development of neurons. They play important roles in metabolism, inflammatory response, neurotransmitter modulation, and synapse function, removal and formation (Barres 2008; Liddelow and Barres 2017; Allen and Lyons 2018). It has been shown in an *in vitro* context that astrocytes are critical to neuronal maturation, synaptically and electrophysiologically (Pfrieger and Barres 1997; Zheng et al. 2025; Hedegaard et al. 2020). Disruptions in astrocyte-neuron crosstalk play a central role in several neurodevelopmental disorders (Sejourne et al., 2024), highlighting the importance of astrocytes in the proper development of neurons.

Metabolic genes are the locus of many genetic differences between humans and our closest non-human primate relatives (Namba et al., 2021; Xing et al., 2024; Stepanova et al., 2021). These metabolic genes play a causative role in neural progenitor cell proliferation. For example, ARHGAP11B, a human-specific gene, increases the amount of basal radial glia through modulating glutaminolysis, the use of glutamine as a carbon source for the TCA cycle (Namba et al., 2020). Additionally, the humanized version of TKTL1 modulates basal radial glia populations by affecting fatty acid synthesis and the pentose phosphate pathway (Pinson et al., 2022).

Studies that compare mice and human cells demonstrate that manipulation of the metabolic features of neural progenitor cells impacts their rate of development. For example, inducing human cells to perform more oxidative phosphorylation, via the inhibition of the enzyme LDHA, leads to an increase in the rate of neuronal maturation (Iwata et al., 2023). Moreover, a preprint study demonstrated that increasing mitochondrial fatty acid oxidation in human organoids drives an increase in maturation rate (Iwata et al., 2025). Collectively, these data suggest that the metabolic properties of brain cells have changed significantly over the course of primate and mammalian evolution and that changes in metabolism can influence the rate of neurodevelopment.

However, these evolutionary studies on cross-species metabolism have primarily focused on basal radial glia or neural progenitor cells. This focus is likely in part because it is proposed that an increase in neural progenitor cell proliferation underlies the unique size and expansion of our cortices in comparison to the cortices of our non-human primate relatives (Kriegstein et al., 2006). However, astrocyte metabolism is critical to neuronal metabolism; astrocytes provide crucial metabolic support to neurons. For example, the astrocyte-neuron lactate shuttle, in which astrocytes produce lactate from glucose and then the lactate is taken up by the neurons for direct entry into the TCA cycle, has been implicated in neuronal activity and long-term memory formation (Suzuki et al., 2011). Inhibition of lactate transporters led to impairments in memory and synaptic plasticity in a mouse model (Suzuki et al., 2011). *In silico* studies have demonstrated that the lactate shuttle leads to much greater theoretical maximum ATP production in neurons (Genc et al., 2011). Furthermore, astrocytes uptake glutamate from neurons to prevent excitotoxicity, and they produce glutamine which can then be used within the astrocyte for TCA cycle entry (Anderson and Swanson 2000).

Finally, astrocytes produce several other building blocks for neurons, such as the amino acids and glia transmitters serine and glycine, fatty acids, and more complex lipids. For example, PHGDH, the enzyme that catalyzes the rate limiting step in serine synthesis, is primarily expressed in astrocytes in the brain (de Koning et al., 2003). Mutations in this gene are associated with microcephaly and cognitive symptoms (Poli et al., 2017; Jaeken et al., 1996; de Koning et al., 2003; Handzlik and Metallo 2023). Collectively, these results demonstrate how many of the metabolic features of neurons are shaped by crosstalk with astrocytes, and many of the functions of neurons are dependent on the metabolites provided by astrocytes.

There is greater transcriptomic divergence between human and non-human primate astrocytes than neurons (Jorstad et al., 2023; Zintel et al., 2024). Human astrocytes, as compared to mouse astrocytes, show lower rates of oxidative phosphorylation and changes in the expression of metabolic genes like *G6PD*, which is higher in mouse astrocytes (Li et al., 2021). These findings collectively suggest evolutionary changes in astrocyte metabolic function. As has been shown in human versus mouse astrocytes, *in vitro* astrocyte studies comparing human, chimp and rhesus astrocytes demonstrated that human versus non-human primate iPSC-derived astrocytes show significant transcriptomic differences in metabolic genes (Ciuba et al., 2025; Zintel et al., 2024). However, despite this evidence of human-specific astrocyte features, research on astrocytes across primate evolution is much more limited than the research on progenitor cells and neurons.

Collectively, these data point to the hypothesis that astrocytes play a crucial role in species-specific neurodevelopment, in part through their role in neuronal metabolism. Our study addresses how astrocytes have changed in primate evolution and whether astrocytes play a role in modulating neurodevelopment in a species-specific manner.

First, we demonstrated successful generation of iPSC-derived astrocytes from several primate species. We also performed a transcriptomic characterization of these astrocytes at three time points during differentiation. Next, we characterized the metabolic properties of these astrocytes to determine if astrocyte metabolism had changed significantly between humans and non-human primates at a functional level. Finally, we tested whether treatment with different species’ astrocyte conditioned media led to differences in neuronal maturation and function. These results were used to assess if astrocytes and more specifically, their metabolic properties, could be contributing to species-specific rates of neuronal maturation and development.

## Results

### Neurons show changes in gene expression associated with metabolic differences in humans versus NHPs

Metabolic genes show significant changes in regulation and coding sequence across the primate lineage, as has been established in the literature (Figure 1A) (Fedrigo et al., 2011; Pinson et al., 2022; Grossman et al., 2004; Burki et al., 2004; Xing et al., 2024; Ju et al., 2025; Stepanova et al., 2021; Florio et al., 2015; Namba et al., 2020; Duka et al., 2014). Re-analysis of sequencing data from Linker et al. (2022) showed differential expression of a key metabolic gene, LDHA, in human as opposed to non-human primate (NHP) neural progenitor cells (NPCs) and neurons, suggesting we were able to recapitulate one of these metabolic differences in our *in vitro* system (Figure 1B). This result was validated by western blot in NPC derived neurons at several time points post-onset of neuronal differentiation (Figure 1C) and has been observed in post-mortem tissue (Duka et al., 2014). To determine whether human and NHP *in vitro* neurons had functional differences in metabolism, we collected NPC-derived neurons five days after the onset of differentiation and sorted them using FACS to obtain a pure neuronal population (Supplemental Figure 1A and 1B, Figure 1D). We then allowed the neurons to recover and performed [U-^13^C_6_] glucose tracing for 24 hours. There were no significant differences in labeling between human and NHP neurons for glycolytic or TCA cycle metabolites (Figure 1E). We also performed tracing with labeled lactate at the same time point in three human and four NHP lines (two gorillas and two chimpanzees). We saw no differences in labeling or abundance in downstream metabolites (Supplemental Figure 1C-H). Intriguingly, a significant difference in the labeling of lactate intracellularly, as well as a trend towards increased labeled abundance (p = 0.06) at the three-week post-differentiation time point was observed, suggesting human neurons took up significantly more lactate than bonobo neurons. However, we did not observe a difference in labeling or abundance in any metabolite downstream of lactate (Supplemental Figure 1I-M). This observation points to the idea that the differences in metabolic gene expression may not be related to the utilization of glucose and lactate but rather to the provision of and initial uptake and usage of lactate by human neurons, although further experiments would be required to conclude this definitively. Interestingly, studies have demonstrated that extracellular lactate changes the expression of LDHA and other glycolytic genes in monocytes, suggesting that LDHA expression may be extrinsically regulated (Schenz et al., 2021). However, it is also possible that the species differences in LDHA expression could be driven by a regulatory change across species, as opposed to external stimuli.

**Figure 1.**
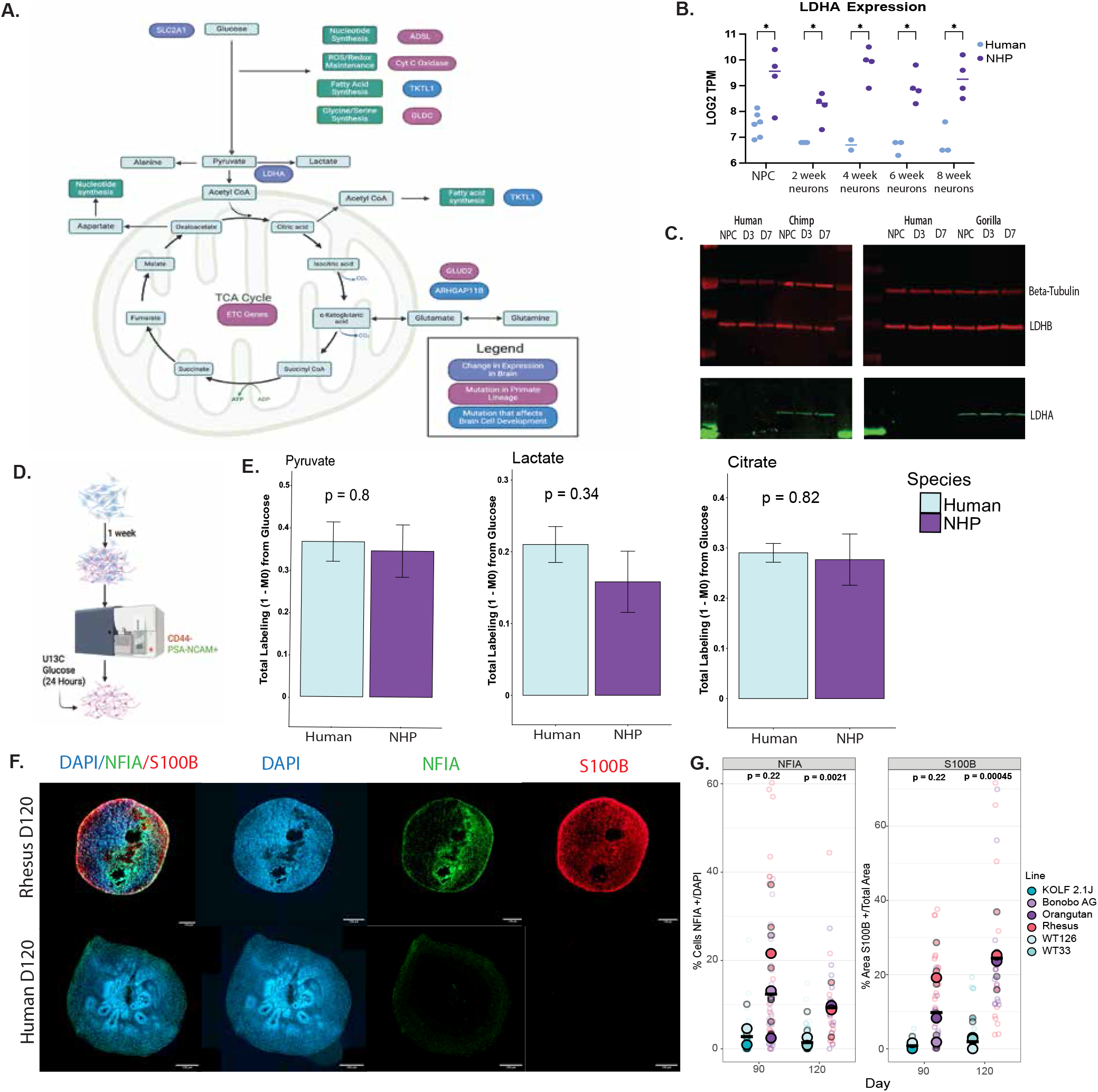
A) Schematic of genes that have changed in human and primate evolution. Genes in purple are those that have established gene expression differences in the brain, those in magenta have a mutation in the primate lineage, and those in blue have a mutation that is known to affect brain development. Made in *Biorender*. B) LDHA expression in NPC and neurons over the course of differentiation, replotted from Linker et al. (2022). Each dot represents a sample. P-value < 0.05 = *. C) Western blot of LDHA protein expression over the course of neuronal differentiation. Cells were collected at the neural progenitor cell (NPC), day 3 post-differentiation, and day 7 post-differentiation. Protein collected from two human lines, one chimp and one gorilla line. Beta-tubulin is a loading control and LDHB is a metabolic gene control. D) Schematic of collection for metabolic data from neurons. E) Human and non-human primate (NHP) tracing data. Technical replicates are averaged together (3 per line). Error bars are standard error of the mean. Y-axis shows what fraction of the given metabolite is labeled from [U-^13^C_6_] glucose. N= 5 cell lines for the NHP group, N = 3 lines for the human group. F) Representative images from 120-day old organoids from human and rhesus macaque. DAPI staining in blue, NFIA in green, S100B in red. Scale bar of length 150 *μ*m. G) Quantification of organoids at day 90 and day 120 for NFIA and S100B. Large circles demonstrate averages per line. Filled in smaller circles indicate organoid averages. Small circles unfilled are individual data points. Quantification done for 4-5 points in each organoid, for 4-5 organoids per line. Humans are in blue, great apes are in purple, and rhesus macaque (old world monkey) are in red. Y-axis indicates the percent of cells (left) or percent of image area (right) positive for the given marker. Statistics performed on line averages.

### Gliogenesis and astrocyte maturation occur earlier in NHP organoids when compared to human organoids

Neurons are dependent on astrocytes for many of their metabolic needs. Therefore, we investigated whether astrocytes may contribute to differences in brain metabolism across primate species. First, we sought to determine whether the timing of astrocyte development and gliogenesis differs across species. We generated forebrain organoids following the protocol in Qian et al. (2018) from three NHP and three human induced pluripotent stem cell (iPSC) lines. We then performed staining for astrocyte markers (early glia marker NFIA and mature marker S100B). Our data demonstrate that astrocytes arise in NHP organoids at earlier time points than in human organoids (Figures 1F and 1G). Given the difference in timing of gliogenesis in three-dimensional cultures, we hypothesized that there may be differences in astrocytic provision of metabolites at early time points in neurodevelopment and that there may be differences in the metabolic role of astrocytes across primate species.

### Astrocytes show differences in metabolic gene expression in humans versus NHPs

Given our hypothesis that metabolic differences between human and NHP astrocytes may exist, we generated a pure *in vitro* system in which we could look specifically at the changes in astrocytes across species. We began by generating glial progenitors and astrocytes from a range of primate species *in vitro* using the protocol described in Santos et al. (2022) and Santos et al. (2017) (Figure 2A). We successfully made astrocytes from three human lines, two chimpanzee lines, two bonobo lines, one gorilla line and one rhesus macaque line. We validated the expression of key glial progenitor and astrocyte markers (Figures 2B and 2C). Furthermore, treatment with IL-1β led to an increase in transcription of several key cytokines and inflammatory responsive genes in both human and NHP astrocytes (Figure 2D), suggesting that our astrocytes had functional immune-responsivity.

**Figure 2.**
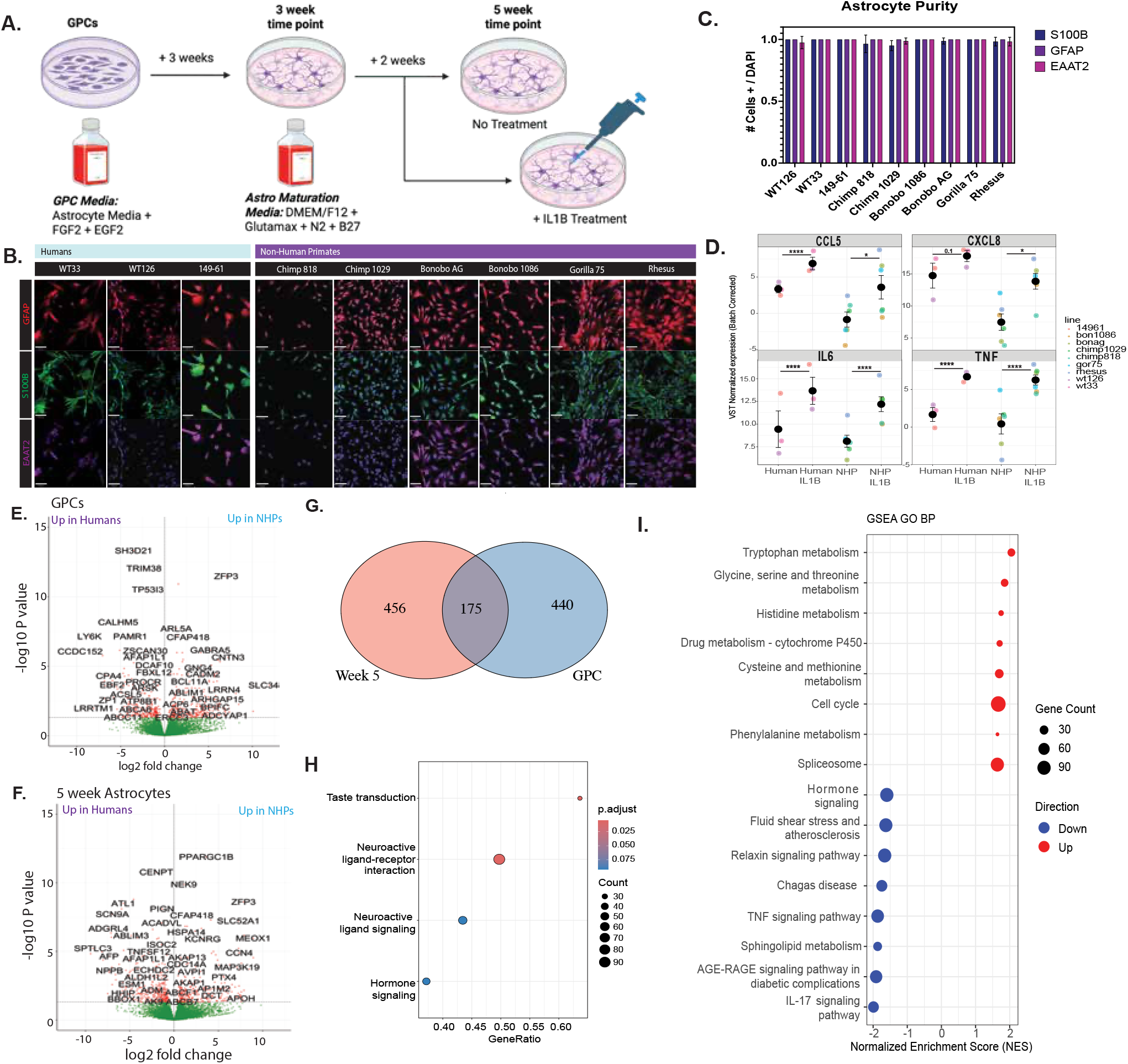
A) Schematic of RNA sequencing collection and differentiation of astrocytes, as well as IL-1β treatment as shown in Figure 2D. B) Staining for key astrocyte markers in all our astrocyte lines. Green shows S100B staining, magenta shows EAAT2 staining, and red shows GFAP staining. C) Quantification of staining as shown in Figure 2B. Each image was divided into quadrants and about 20 cells were counted for the presence or absence of the given marker. Y-axis shows the number of cells that were positive/DAPI averaged over the four quadrants. D) Plot showing the changes in expression of key cytokines and inflammatory-responsive genes in response to IL-1β treatment. Four key genes are shown. Each data point is the average of technical replicates for each line, and dot color indicates line. Statistics were performed by comparing human to human + IL-1β and NHP astrocytes versus NHP astrocytes + IL-1β. Black circles indicate mean, and error bars indicate standard error. P-values are adjusted p-values from DESEQ2 with the formula ∼ line + IL-1β E) Volcano plot of differentially expressed genes between human and NHP genes at the GPC time point. Positive LFC indicates increased expression in NHPs, negative LFC indicates increased expression in humans. F) Volcano plot of differentially expressed genes between human and NHP genes at the five-week-old differentiated astrocyte time point. G) Venn diagram showing differentially expressed genes between human and NHPs at the GPC stage (right) and five-week-old astrocytes (left). 175 genes are differentially expressed at both stages of maturation. H) GSEA for GPC RNA-sequencing data comparing human and NHP GPCs. Color shows p-value. Size of dot shows how many genes drive enrichment in the given category. I) GSEA of five-week-old astrocyte RNA-sequencing data comparing human and NHP data in DESEQ2. Blue (negative normalized enrichment score) indicates higher expression of the pathway in human astrocytes. Red indicates higher expression of the pathway in NHP astrocytes. Dot size indicates how many genes drive enrichment in the given category.

To understand the differences in human versus NHP astrocytes, we began with a transcriptomic characterization of our human and NHP astrocytes at the glial progenitor cell (GPC) stage, three weeks and five weeks into *in vitro* maturation (Supplemental Figure 2A). We decided to focus on the GPC and five-week post differentiation stages for further downstream analyses. We ran differential expression analysis to determine which genes were differentially expressed between our human and NHP cell lines at each time point. We identified 615 differentially expressed genes at the glial progenitor stage and 631 differentially expressed genes at the five-week-old astrocyte state (p adjusted < 0.05) (Figure 2E-G). There were 175 genes that were significantly differentially expressed at both the GPC and five-week-old time point, and enrichment analysis primarily showed these shared differentially expressed genes were related to neuronal signaling (Figure 2G, Supplemental Figure 2B). Furthermore, there was a significant correlation between log fold change NHP/human as measured by DESEQ2 at the GPC state and astrocyte state, suggesting there were conserved differences in gene expression across species regardless of differentiation stage (Supplemental Figure 2C).

Gene set enrichment analysis showed an enrichment for genes related to neuronal signaling and ligands at the GPC state (Figure 2H). At the five-week time point, we observed an upregulation of genes involved in immune response in the human astrocytes (Figure 2I, Supplemental Figure 2D). This trend in our data corresponds to previous literature that human astrocytes have a stronger response to inflammatory stimuli than mouse astrocytes (Li et al., 2021). Gene set enrichment analysis showed an enrichment for the upregulation of metabolism associated pathways in our NHP astrocytes at the five-week time point (Figure 2I, Supplemental Figure 2D). We specifically observed an enrichment for amino acid metabolism (tryptophan metabolism, glycine, serine and threonine metabolism, histidine metabolism, phenylalanine metabolism, and cysteine and methionine metabolism are all enriched pathways). These results further support the changes in metabolism genes identified in Ciuba et al. (2025) when comparing human and chimpanzee *in vitro* astrocytes. Furthermore, when we compared the expression of key enzymes involved in glycolysis and glycine and serine metabolism (two pathways that were enriched), we observed a general upregulation of glycine and serine metabolism pathway enzymes in the NHP astrocytes. We also observed a general downregulation of glycolysis genes (most notably *HIF1A* and *SLC16A3*, a key aerobic glycolysis and lactate transport gene, respectively) in the NHP astrocytes compared to the human astrocytes at the GPC state and the five-week-old astrocyte time point (Figure 3A).

**Figure 3.**
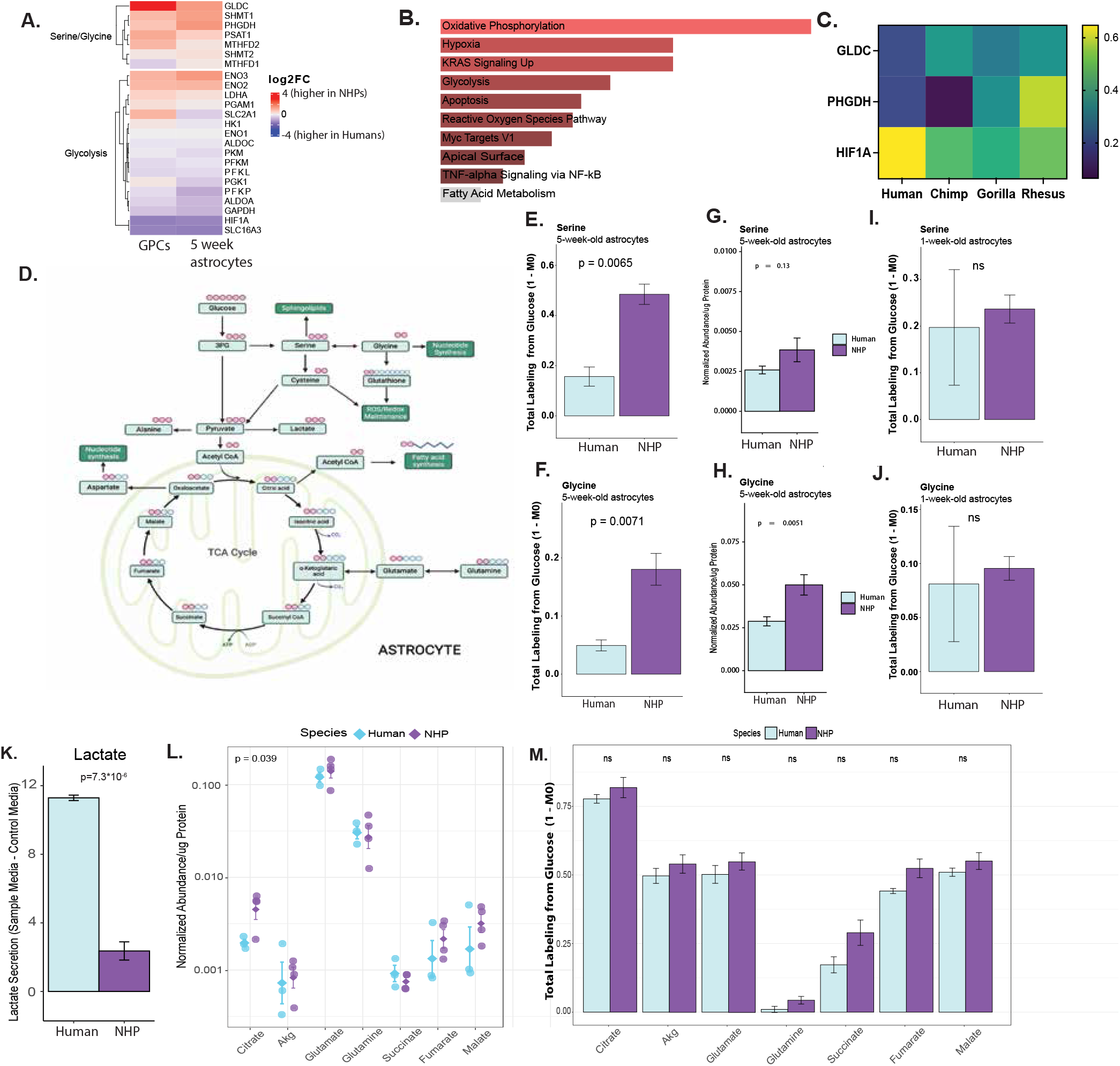
A) Expression of key metabolic genes at the GPC and five-week time points. Color indicates LOG2FC, with a positive log2 fold change indicating higher expression in NHP astrocytes compared to human astrocytes and a negative value indicating higher expression in human astrocytes. LFC values generated with DESEQ2. B) MSigDB Hallmark enrichment for genes differentially expressed between human and chimpanzee astrocyte clusters in post-mortem data. Differential expression values and data used from Jorstad et al. (2023). Analysis done with Enrichr using MSigDB Hallmark 2020 Analysis. Red coloring indicates unadjusted p-value < 0.05. C) Average expression of a few key genes in astrocyte clusters from sn-RNAseq data in different species (re-graphed from Jorstad et al., 2023 and Cytosplore Viewer). D) Schematic of [U-^13^C_6_] glucose carbon labeling experiments. Red and blue circles indicate carbons. Red carbons indicate those that may end up labeled by the addition of [U-^13^C_6_] glucose. E) Serine labeling from [U-^13^C_6_] glucose in 5-week-old astrocytes. Y axis shows 1 minus the unlabeled amount over the total amount of the given metabolite. N = 2 for human and N=5 for NHP (biological replicates). Technical replicates are averaged together. Bars show standard error. F) Glycine labeling from [U-^13^C_6_] glucose trace in five-week-old astrocytes. G) Intracellular serine abundance normalized to protein amount and an internal standard in five-week old astrocytes. H) Intracellular glycine abundance normalized to protein amount and an internal standard in five-week old astrocytes. I) Serine labeling from [U-^13^C_6_] glucose trace in one-week-old astrocytes. J) Glycine labeling from [U-^13^C_6_] glucose trace in one-week-old astrocytes. K) Lactate abundance in media in human versus NHP astrocyte samples. Bars denote standard error. Values normalized to baseline media (unconditioned) abundances. Technical replicates run in triplicate and averaged together per biological replicate (line). Biological replicates: n = 2 for human and n=6 for NHPs. Bars show standard error. L) Normalized abundance of TCA cycle intermediates and offshoot metabolites measured intracellularly using a GC-MS. Values normalized to norvaline, an internal control, and the amount of protein. Technical replicates averaged together; biological replicates are individual lines. N = 2 for human and N=5 for NHPs. Y-axis is log scale. Bars show standard error. M) TCA cycle labeling from [U-^13^C_6_] glucose in 5-week-old astrocytes. Y axis shows 1 minus the unlabeled amount over the total amount of the given metabolite. N = 2 for human and N = 5 for NHP (biological replicates). P <0.05 is labeled. Bars show standard error.

Because an *in vitro* setting is metabolically different than the brain, and because our *in vitro* astrocytes are immature compared to the astrocytes in the brain, we turned to post-mortem single nuclei transcriptomics data to validate the enrichment for metabolic genes in human versus chimpanzee astrocyte gene expression. Jorstad et al (2023) performed post-mortem single nuclei transcriptomics on human, chimpanzee, gorilla, rhesus and marmoset middle temporal gyrus. We extracted the differentially expressed genes in the astrocyte cluster from their analysis comparing human and chimpanzee astrocytes. The top pathways in MSigDB enrichment analysis (ranked by p-value) included oxidative phosphorylation, hypoxia (related to *HIF1A*), and glycolysis (Figure 3B). Furthermore, we looked at the average expression of *GLDC*, *PHGDH*, and *HIF1A* (three key metabolic genes) in the astrocyte clusters in this data. *GLDC* and *PHGDH* both encode enzymes critical to serine and glycine metabolism, and *HIF1A* is a transcription factor that regulates aerobic glycolysis. We observed a trend towards decreased expression of *PHGDH* and *GLDC* in human compared to chimp, gorilla and rhesus astrocytes in post-mortem tissue, and an increase in *HIF1A* expression in humans, which parallels what we observed *in vitro* (Figure 3C). However, we were unable to perform statistics comparing these values because we used the average expression in each species cluster as reported in Cytosplore Viewer, and therefore, there is only one value for the human samples. These expression differences indicate that these metabolic gene expression differences *in vitro* may also be relevant to astrocytes in the adult brains of humans and NHPs.

### NHP astrocytes show widespread metabolic restructuring compared to human astrocytes

Next, we investigated whether the significant differences we observed in astrocyte metabolic gene expression translated to functional species differences in astrocyte metabolism. To quantify and compare metabolite production between humans and other primates, we performed [U-^13^C_6_] glucose labeling experiments in our human and NHP lines at the one week and five-week time points (Figure 3D). At the five week time point, we observed an increase in intracellular glycine and serine labeling from glucose in the NHP astrocytes (5 lines; one bonobo, two chimp, one gorilla, one rhesus) compared to the human astrocytes (2 lines) (Figure 3E and F; Supplemental Figure 3A-F). We also observed a significant increase in glycine abundance as well as a trend towards increased serine abundance in the NHP astrocytes (Figure 3G and 3H; Supplemental Figure 3G-I). We interpreted these data as demonstrating an increase in *de-novo* synthesis from glucose of serine and glycine in the NHP astrocytes compared to the human astrocytes. This finding aligned with data from our transcriptomics assay in five-week-old astrocytes, which showed an upregulation of serine, glycine and threonine metabolism in NHP astrocytes (Figures 2I, 3A and 3C). This difference in serine and glycine labeling was not observed at our one-week time point, suggesting that this phenotype is found at more mature time points in astrocyte differentiation (Figures 3I and 3J, Supplemental Figure 3J-3L). This assay at the one-week time point was performed in two human lines, two chimp lines, one bonobo line, one gorilla line, and one rhesus line.

Human astrocytes showed a substantial increase in lactate release into the media as opposed to NHP astrocytes (Figure 3K). This observation correlated with our transcriptomic findings, which demonstrated an increase in the expression of *HIF1A* and *SLC16A3* (the main lactate exporter in astrocytes, MCT4) in human astrocytes.

We also observed an upregulation of intracellular citrate abundance but not succinate or other downstream TCA cycle metabolites (Figure 3L) in NHPs but no differences in labeling (Figure 3M). An increase in citrate abundance without a corresponding increase in other TCA metabolites could be consistent with increased citrate availability for acetyl-CoA production, prompting us to investigate downstream fatty acid labeling (Icard et al., 2021). We observed a trend towards an increase in *de novo* synthesis of palmitate and oleic acid from glucose (Supplemental Figure 3M and 3N). However, acetyl-CoA could also be used for acetylation, play a role in epigenetics, be used for the synthesis of other lipids such as cholesterol, or serve as a signaling molecule in other cellular pathways (Westergaard et al., 2017; Icard et al., 2021). In general, an increase in citrate abundance is associated with a reduction in aerobic glycolysis, as citrate inhibits PFK (Garland et al., 1963; Newsholme et al., 1977). Since an increase in glucose oxidation to citrate may also suggest an overall increased metabolic rate, we sought to evaluate other TCA cycle related metabolites. We did not observe a significant increase in label or abundance of other TCA cycle related metabolites (Figure 3M), further supporting our hypothesis that NHP astrocytes had increased flux through anabolic pathways, supporting the synthesis of key metabolic precursors, as opposed to merely operating at a higher metabolic rate.

Given that we did not observe a significant difference in citrate labeling from glucose (Figure 3M), we tested whether a different carbon source was generating the increase in citrate abundance we observed. To test the hypothesis that the NHP astrocytes were using more glutamine as opposed to glucose as a carbon source, we performed a tracing experiment on our cells at the five-week time point using [U-^13^C_5_] glutamine. We did not observe a significant different in glutamine enrichment in TCA cycle intermediates, although there was a trend towards increased labeling in the NHPs, suggesting that increased glutamine utilization might contribute to the increase in citrate abundance, but it is likely not the main explanatory reason in our system (Supplemental Figure 3O).

Collectively, these data suggested an astrocytic metabolic reprogramming in humans after they diverged evolutionarily from NHPs. Human astrocytes primarily generate and release lactate whereas the NHP astrocytes perform more anabolism of key building blocks, which may be important for neuronal differentiation and maturation.

### NHP astrocyte conditioned media leads to changes in expression of neuronal differentiation-related genes and faster electrophysiological maturation in human neurons

Given the significant differences in metabolism we observed in the human and NHP astrocytes, we investigated whether there is a differential impact of human versus NHP astrocyte metabolism on neuronal functioning and maturation, as there is a well-established observation in the literature that human cortical neurons undergo a more prolonged developmental and maturation program than those of NHPs. (Marchetto et al., 2019; Somel et al., 2009; Schörnig et al., 2021; Pollen et al., 2019; Vanderhaeghen and Polleux 2023). We hypothesized that the NHP astrocytes, through their provision of key metabolic intermediates that support neuronal growth and differentiation, may serve as a cell-extrinsic mechanism for changing the rate of neuronal maturation and differentiation trajectories.

To test this hypothesis, we cultured astrocytes from three human lines and four NHP lines (one chimp, one bonobo, one gorilla and one rhesus line) in neuronal media for twenty-four hours. We collected that media and fed it to human neural progenitor cells (NPCs), and we measured changes in their transcriptome and in their electrical activity (Figure 4A). For our transcriptomics study, we fed the NPCs astrocyte conditioned differentiation media for 1 day, 4 days, 7 days or 14 days. We then performed differential expression analysis between the human and NHP astrocyte conditioned media treated neurons and performed GO analysis. We observed an enrichment for terms related to neuronal differentiation, neuron fate commitment, and forebrain development, suggesting that NHP astrocyte conditioned media with human neurons causes changes in the trajectory of neuronal maturation and differentiation (Figure 4B). To further investigate these transcriptomic changes, we looked at the genes that were differentially expressed in three or more of the time points studied (Figure 4C and 4D). We identified ten genes that showed significant differential expression between human and NHP conditioned media conditions in at least three of the four time points (Figure 4D). We observed an upregulation of *SMAD6* in neurons exposed to NHP astrocyte conditioned media and an upregulation of *EBF2*, *VXN*, *NEUROD4*, *NEUROD1*, *NEUROG1* and *NEUROG2* in neurons exposed to human astrocyte conditioned media. *SMAD6* is known to promote neuronal differentiation (Xie et al., 2011). *EBF2* is a marker of newly committed neurons (Chuang et al., 2011; Garcia-Dominguez et al., 2003), and both *NEUROG1* and *NEUROG2* coordinate cell cycle exit and neuronal differentiation by functioning as early pioneer transcription factors that establish the neuronal gene regulatory program (Lacomme et al, 2012; Bertrand et al., 2002). *VXN* is transiently expressed at the beginning of neuronal differentiation in progenitors early in cell fate commitment (Moore et al., 2018). These genes which are upregulated in the human astrocyte conditioned media treated neurons are early markers of neuronal cell fate commitment and help establish and maintain developmental transcriptional programs rather than marking terminal neuronal maturation. We hypothesize that their sustained expression may therefore reflect a more prolonged developmental state, consistent with the delayed maturation observed in human neurons in previous studies. Collectively, these data demonstrate a change in neuronal differentiation trajectories in those human neurons cultured with NHP versus human astrocyte conditioned media and a change in the expression of key regulators of neuronal differentiation.

**Figure 4.**
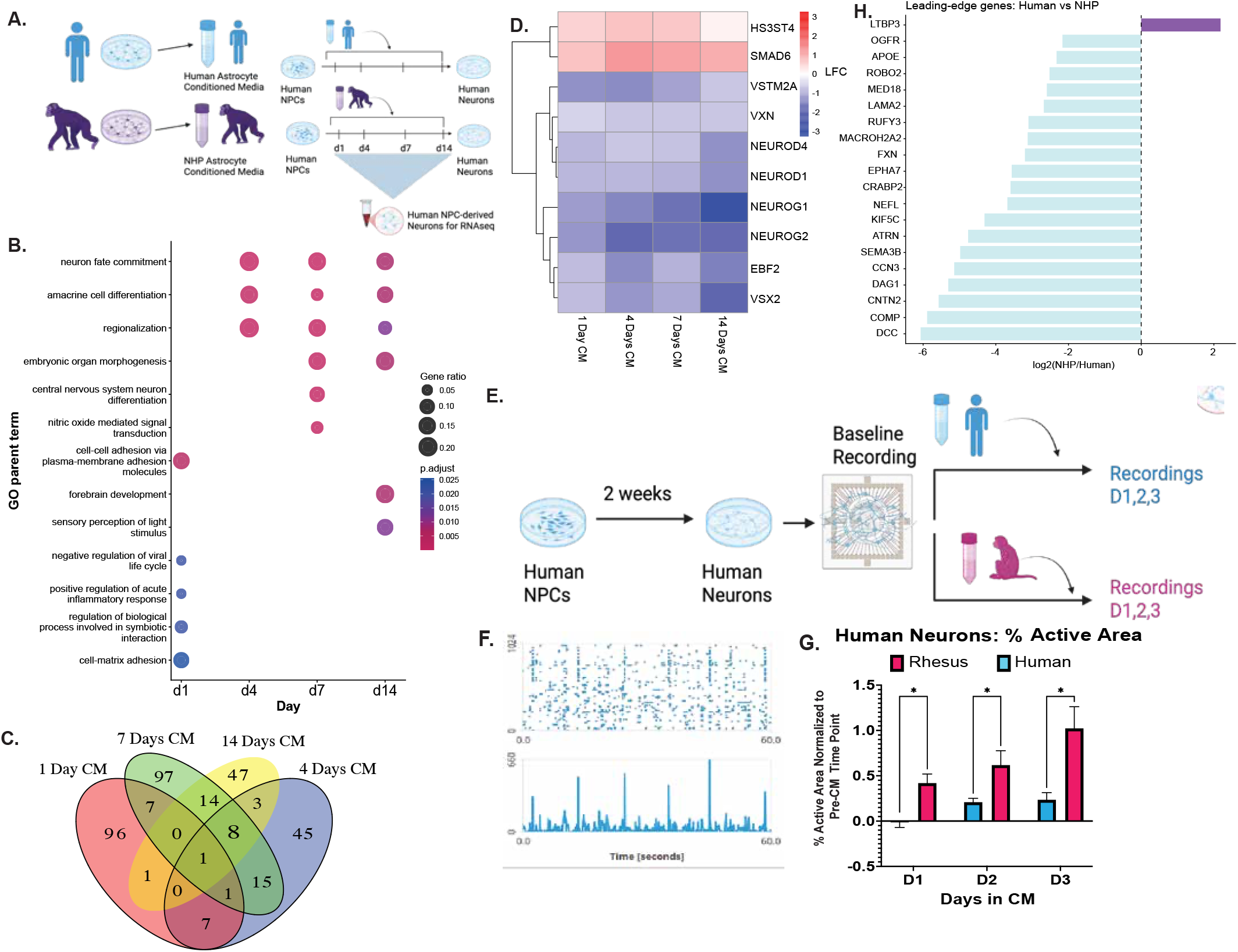
A) Schematic of conditioned media experiments. Neuronal differentiation media was added to human or NHP astrocytes and then collected, filtered, and added to human NPCs. Three human astrocyte lines were used and four NHP lines were used. B) GO term analysis from RNA-sequencing from human neurons cultured with human versus NHP astrocyte conditioned media. Dot colors indicate adjusted p-value for enrichment. Gene ratio indicates the number of genes in the given category that are significantly differentially expressed. X-axis indicates the days in conditioned media before the NPC-derived neurons were collected for RNA. C) Venn diagram of differentially expressed genes (FDR < 0.1) in neurons between human and NHP astrocyte conditioned media treatment for each time point (duration in conditioned media). D) Genes that were differentially expressed (FDR <0.1) in three out of four of the conditions were plotted. Color indicates log fold change between human conditioned media neurons and NHP conditioned media neurons. Red indicates higher expression of the given gene in NHP astrocyte conditioned media condition, blue indicates higher expression in human astrocyte conditioned media condition. LFC shown for each duration of time in conditioned media (1 day in CM, 4 days, 7 days, or 14 days). E) Leading edge proteins from secretome of human versus NHPs for top GSEA categories. Astrocyte secreted proteins more abundant in human samples are in light blue, and those more abundant in NHP samples are in purple. N = 3 lines (2 technical replicates per line) for human and 4 for NHP lines (2 technical replicates per line). F) Schematic for MEA experiments. Human and rhesus astrocyte conditioned media collected and added to two-week-old NPC-derived neurons seeded onto an MEA plate. Recordings normalized to baseline recording prior to addition of any astrocyte conditioned media. G) Representative trace of firing activity for multi-electrode array assay. In the upper plot channels, the y-axis indicates different channels of electrical activity recorded, and the x-axis represents the time in the recording (from 0 to 60 seconds). The bottom plot indicates a combined network activity score as produced by the Maxwell software for each of the time points. H) Plot of the percent area of a well with activity during an activity scan. Each well was normalized to its own pre-conditioned media recording time point. Blue bars indicate wells of human neurons fed human astrocyte conditioned media. Red indicates wells of human neurons fed rhesus astrocyte conditioned media. P-values indicate differences between human and rhesus activity at each time point. X-axis shows days in conditioned media. Neurons were differentiated for two weeks and replated on MEA plate before the addition of species-specific conditioned media. N = 5 for human conditioned media and N=5 for rhesus. Error bars show standard error. Asterisks indicate p < 0.05.

To assess the role of astrocytes in modulating species-specific rates of neuronal maturation and development, we tested whether neurons exposed to conditioned media showed differences in electrical activity, a canonical metric of neuronal maturation. We treated two-week-old neurons from humans with conditioned media from one of two human lines or from one rhesus line for a total of three days (Figure 4E). Relative to the electrical activity in each well prior to the addition of astrocyte conditioned media, there was an increase in the percent of active electrodes in those human neurons exposed to rhesus conditioned media as opposed to human astrocyte conditioned media (Figure 4F and 4G). This result suggested an increase in electrophysiological maturation in those human neurons exposed to rhesus astrocyte conditioned media. We interpreted this as the faster achievement of a terminal maturation state in the NHP conditioned media neurons, which again fits with the neoteny phenotype previously observed by others when comparing human and NHP neurons.

In order to understand how the astrocytes may be playing a role in these differing neuronal trajectories, we analyzed the proteomic secretome of human and NHP astrocytes. Using four NHP lines (one bonobo, one chimpanzee, one gorilla, and one rhesus line) and three human lines, we performed proteomics on the astrocyte conditioned media. When we performed principal component analysis on these secretomic data, we saw a strong divide along principal component two between human and NHP samples, suggesting that the secretome was divergent between species groups (Supplemental Figure 4A). When we looked at the differentially secreted proteins using MSstats, we saw similar numbers of proteins upregulated in both primate and human astrocyte conditioned media (Supplemental Figure 4B). Examining the top differentially secreted proteins by p-value (Supplemental Figure 4C) revealed that several key synaptogenic or neuronal maturation related proteins such as *APOE* (Tensaouti et al., 2018; Yu et al., 2021), *NRCAM* (Sakurai et al, 2012) and *LAMA2* (Arreguin et al., 2020; Anderson et al., 2005) were secreted at significantly higher levels by the human astrocytes. Running GSEA GO on the secretomics data set supported this observation, as this analysis revealed an upregulation in human astrocyte conditioned media of a number of terms associated with synaptogenesis and neuronal branching, such as axon development and neuron projection guidance/development (Supplemental Figure 4D). The leading edge genes from this GSEA revealed that several key genes involved in neuronal branching and axon development/guidance, such as *ROBO2* (Blockus et al., 2021; Zhang et al., 2012; Campbell et al., 2007), *SEMA3B* (Mohan et al., 2019), *DCC* (Horn et al., 2013), *EPHA7* (Clifford et al., 2014), and *LAMA2* (Arreguin et al., 2020), are found at higher amounts in human astrocyte conditioned media (Figure 4H). These proteins are not necessarily associated with a faster developmental trajectory in neurons, but instead are associated with synaptogenesis and neuronal complexity/dendritic arborization. This suggested that the human astrocytes were more involved in shaping synaptic signaling and dendritic arborization than the NHP astrocytes.

Taken together, our transcriptomic, electrophysiological, and secretomic data support a model in which human and NHP astrocytes shape neurodevelopment trajectories differently. While NHP astrocytes appear to promote more rapid achievement of a terminal maturation state, human astrocytes preferentially provide cues associated with prolonged neuronal development transcriptional programs, synaptic development and neuronal structural complexity, suggesting that species-specific astrocyte programs contribute to the distinct developmental trajectories of NHP versus human neurons.

### PHGDH and TGFβ1 can recapitulate the metabolic restructuring of human astrocytes in NHP astrocytes

We investigated broader cellular pathways that may regulate the differences in metabolism we observed. We identified a general upregulation of TGFβ signaling pathways in human astrocytes in our transcriptomics dataset (Supplemental Figure 5A). TGFβ has been previously implicated as one of the regulators of metabolism (Liu et al., 2022). To identify whether TGFβ may be upstream of the differences in metabolism we observed, we added TGFβ1 to the media of astrocytes from one human line and from one NHP (chimpanzee) line. We then performed a [U-^13^C_6_] glucose trace. In both the human and NHP lines, we observed a significant decrease in *de novo* serine and glycine synthesis (Supplemental Figure 5B and 5C). We also observed an increase in lactate secretion in the chimpanzee and human astrocytes supplemented with TGFβ1, indicating that TGFβ1 might also lie upstream of the differences in lactate processing we observed (Supplemental Figure 5D). We observed a decrease in intracellular serine and glycine abundance in both species (although glycine abundance in human astrocytes was not significantly lower with TGFB1 treatment) (Supplemental Figure 5E and F).

Given the differences in serine synthesis we had observed, we also investigated whether we could recapitulate our metabolic phenotype by inhibiting the rate limiting enzyme in serine synthesis, PHGDH, using the small molecule NCT-503. We used one chimpanzee line and one human line and added NCT-503 while performing a [U-^13^C_6_] glucose trace. We observed a decrease in *de novo* serine and glycine synthesis (Figures 5A and B) in both the human and chimpanzee cells, but the decrease was notably more severe in the chimpanzee cells, suggesting that the chimpanzee cells are more sensitive to PHGDH inhibition. We also observed a significant decrease in serine and glycine abundance in both the human and chimpanzee line with PHGDH inhibition (Supplemental Figure 5G and H). Furthermore, we observed that inhibiting PHGDH led to an increase in lactate release from the astrocytes in both the chimpanzee and human astrocytes, again with a more notable increase in the chimpanzee astrocytes (Figure 5C). This result was intriguing, as PHGDH is specifically involved with serine synthesis but NCT-503 is modulating lactate release as well in our system. Collectively, these data suggest we can recapitulate many of the metabolic features of human astrocytes by inhibiting PHGDH in chimpanzee astrocytes.

**Figure 5.**
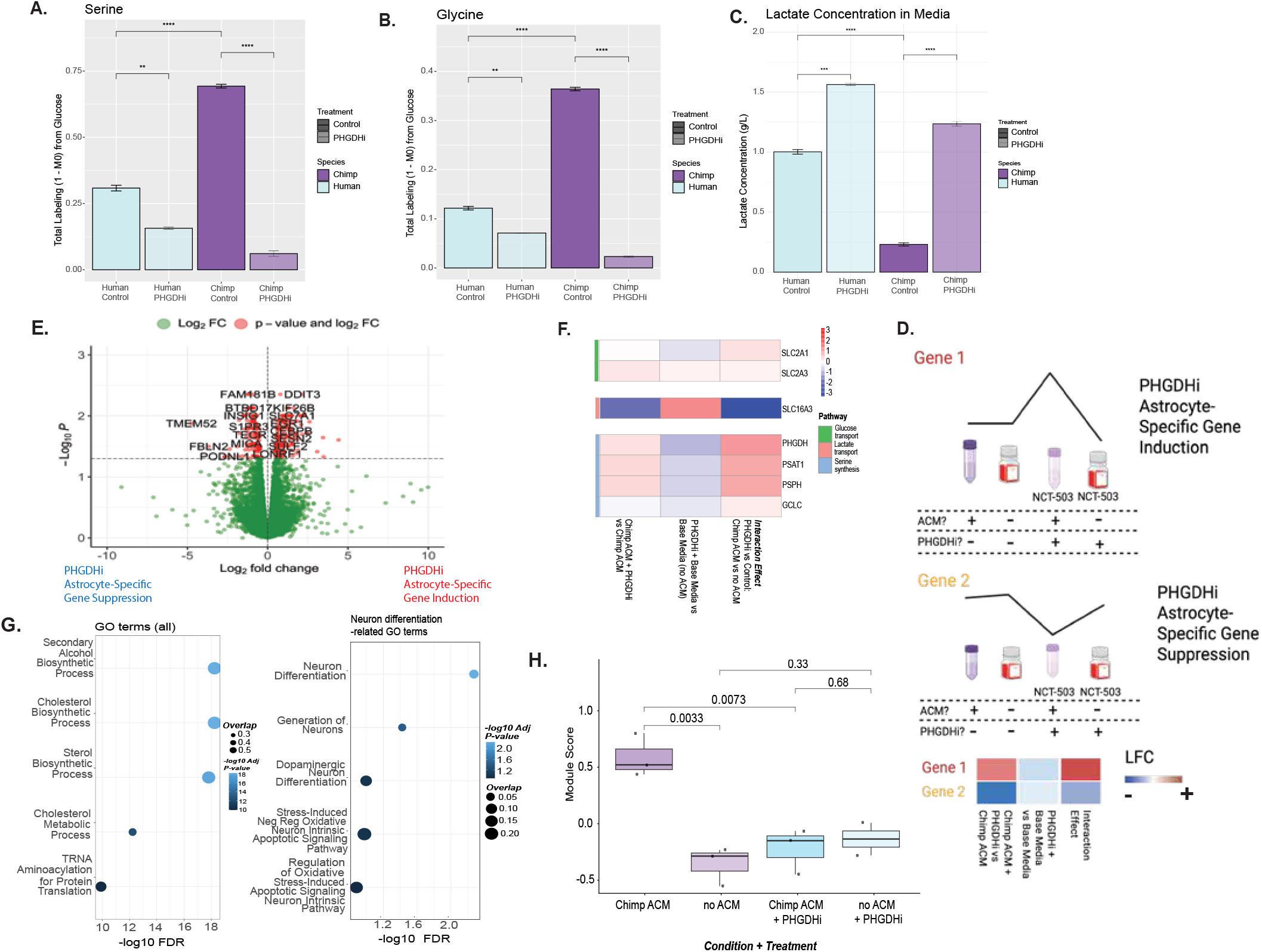
A) Serine labeling from [U-^13^C_6_] glucose in 5-week-old astrocytes treated with a PHGDH inhibitor. Y axis shows 1 minus the unlabeled amount over the total amount of the given metabolite. Bars show standard error. * = p < 0.05; ** = p < 0.01 ; *** = p < 0.001 ; **** = p < 0.0001. Purple indicates chimpanzee astrocyte media, and light purple shows media from chimp astrocytes treated with a PHGDH inhibitor (NCT-503) for 24 hours. Blue indicates human astrocyte media, and lighter blue shows media from human astrocytes treated with a PHGDH inhibitor. B) As described for Figure 5A but for glycine. C) Lactate concentration in media as measured by YSI. Purple indicates chimpanzee astrocyte media, and light purple shows media from chimp astrocytes treated with a PHGDH inhibitor (NCT-503). Blue indicates human astrocyte media, and lighter blue shows media from human astrocytes treated with a PHGDH inhibitor. One chimpanzee line and one human line were used, and different wells were treated as replicates. * = p < 0.05; ** = p < 0.01 ; *** = p < 0.001 ; **** = p < 0.0001. D) Schematic of PHGDH inhibition RNA-sequencing experiments. Representative heatmap is shown below (not real data). E) Volcano plot of genes differentially regulated in response to PHGDH inhibition in chimpanzee astrocytes. DESEQ2 formula of treatment + conditioned media + treatment: conditioned media was used. Positive LFC indicates genes that show increases in gene expression specifically due to the effect of PHGDH inhibition on the chimp astrocytes. Negative LFC values indicate genes that are downregulated. F) Key metabolic gene expression in neurons in response to PHGDHi in chimp astrocytes. LFC is indicated in the colors. On the x-axis, LFC is shown for, respectively, comparing the expression in neurons exposed to chimp astrocyte conditioned media with PHGDH inhibition vs regular chimp astrocyte conditioned media; just the PHGDH inhibitor in the base neuronal media versus base neuronal media without inhibitor; and the interaction effect of PHGDH inhibition with the astrocyte conditioned media. Key genes were picked based on their involvement in metabolic pathways that changed in astrocytes (see Figure 3). G) GO analysis of genes differentially regulated due to the effect of PHGDH inhibition on the chimp astrocytes. DESEQ2 formula of treatment + conditioned media + treatment: conditioned media was used. Left side shows top enriched GO terms overall for differentially expressed genes, with size of the dot indicating the amount of overlap with all the terms in the category. X-axis shows -log10 of FDR and color of dot indicates -log10 p-value. Top 5 terms are shown. The graph on the right shows the top significantly enriched GO terms related to neuronal functioning. H) Module score for differentially expressed genes between human and NHP media conditions at one time point (the day four time point) when applied to the PHGDH inhibitor experiment. X-axis shows neurons in different conditions (chimp astrocyte conditioned media, regular base media, chimp astrocyte conditioned media from chimp astrocytes treated with PHGDH inhibitor, and regular base media with PHGDH inhibitor).

### PHGDH inhibition in astrocytes alters the transcription of neuronal differentiation genes in neurons cultured with astrocyte conditioned media

Next, we tested how these differences in metabolism affected neurons cultured in astrocyte conditioned media after the astrocytes had been treated with the PHGDH inhibitor. Specifically, we were interested in whether we could recapitulate the changes observed in neuron differentiation by specifically manipulating the metabolism of the astrocytes through PHGDH inhibition. We used four conditions: chimp astrocyte conditioned media, chimp astrocyte conditioned media + PHGDH inhibitor, base neuronal differentiation media, and base neuronal differentiation media + PHGDH inhibitor. Using DESEQ2, we looked for genes that showed an interaction effect for PHGDH inhibition and astrocyte conditioned media. This approach allowed us to identify genes that responded in neurons specifically to the impact of PHGDH inhibition on the astrocytes, not just the genes that responded to the presence of the PHGDH inhibitor in the media (Figure 5D). Genes that showed a positive log fold change were identified as genes with PHGDHi-astrocyte-specific gene induction, meaning that they were upregulated in response to both the presence of PHGDH inhibitor and astrocyte conditioned media but not just one or the other. Those genes downregulated in this analysis were identified as PHGDHi astrocyte-specific suppressed genes (Figure 5D).

When we analyzed the data we identified 301 differentially expressed genes (adjusted p-value < 0.05) (Figure 5E). When we inhibited PHGDH in the astrocytes, we observed an upregulation of several serine and glycine related genes in the neurons specifically in the interaction analysis, suggesting that the neurons had to make their own serine and glycine without provision by astrocytes. *SLC16A3*, which is a gene related to lactate export, was also suppressed in the interaction effect, which may be because the astrocytes were producing much more lactate and thus the neurons may not have been synthesizing and exporting as much lactate (Figure 5F). Collectively, this result indicated that there was a remodeling of neuronal metabolic transcription programs in response to changes in the metabolism of astrocytes.

When we performed GO enrichment on the differentially expressed genes, many gene sets associated with metabolism were significantly enriched (Figure 5G), as expected. Several GO terms related to neuronal functioning were also significantly enriched, and the top two GO terms were neuron differentiation and generation of neurons (Figure 5G).

We next tested whether the changes in genes that were differentially expressed in response to astrocyte conditioned media from NHPs versus humans were mediated by PHGDH expression in the astrocytes. We created a module score of the genes significantly upregulated in NHP astrocyte conditioned media versus human astrocyte conditioned media at the four day time point. We observed higher expression (as indicated by a higher module score) of those genes in the chimpanzee astrocyte conditioned media versus the no astrocyte conditioned media condition, as expected. PHGDH inhibition in the chimpanzee astrocytes led to expression levels similar to the levels in neurons grown without astrocyte conditioned media (Figure 5H). This result indicates that PHGDH inhibition in the astrocytes may ameliorate the effect of chimp astrocyte conditioned media on neuronal gene expression and that the metabolic rearrangement of NHP astrocytes may contribute to or be necessary to the change in neuronal development trajectory driven by NHP astrocytes.

## Discussion

Neuronal development, function, and maintenance are highly dependent on astrocytes; astrocytes assist with synaptic pruning, electrophysiological maturation, metabolism, and immunological responses (Barres et al., 2008). Therefore, we investigated whether there are species-specific differences in astrocytic function that may influence species-specific features of neuronal development.

In this study, we compared human and NHP *in vitro* astrocytes and examined their role in neuronal functioning and maturation. We demonstrated successful generation of iPS-derived astrocytes from six NHP individuals, across four species, allowing us to study astrocyte function in a diverse cohort of NHPs. We also demonstrated a decrease in the expression of metabolic genes, mostly related to amino acid anabolism, in human astrocytes in comparison to NHP astrocytes. We revealed a human-specific specialization in lactate production, whereas the NHP astrocytes were characterized by an increase in serine and glycine synthesis. Critically, we demonstrated an increase in electrophysiological maturation and changes in the expression of neuronal differentiation genes in neurons exposed to astrocyte conditioned media from NHPs as compared to humans.

Comparisons between mouse and human cells have previously identified changes in metabolic function. Li et al. (2021) demonstrated a decreased OCR/ECAR ratio in human astrocytes compared to mouse astrocytes, suggesting that human astrocytes show an increased reliance on glycolysis. In this study, mouse astrocytes performed more pentose phosphate pathway metabolism and oxidative phosphorylation. Furthermore, a comparison of mouse and human immune cells showed significant changes in serine metabolism in activated NK cells (Li et al., 2025). However, to our knowledge, functional metabolic studies have not been done in human versus NHP astrocytes.

In our data, we observed a human-specific increase in lactate secretion in astrocytes. These data could indicate a species-specific divergence in the astrocyte-neuron lactate shuttle. Neurons can use the lactate as an energy source feeding directly into the TCA cycle (Karagiannis et al., 2021; Wyss et al., 2011). *In silico* studies have shown that lactate provision by astrocytes increases the maximum amount of ATP a neuron can make, which could help support a more energetically demanding neuron (Genc et al., 2011). In turn, human neurons have been shown to be larger and more morphologically complex cells (Libe-Phillipot et al., 2024; Lindhout et al., 2024) that may require more ATP to support action potentials (Sengupta et al., 2013). However, the increases in serine and glycine synthesis and citrate abundance suggest that NHP astrocytes are biosynthetically more active, potentially synthesizing more proteins or lipids, which may be helpful in providing building blocks for developing and maturing neurons.

Mutations in PHGDH, the rate limiting enzyme in serine synthesis, can lead to neurological syndromes and changes in neuronal differentiation/morphology and expansion (Poli et al., 2017; Jaeken et al., 1996; de Koning et al., 2003; Handzlik and Metallo 2023). PHGDH haploinsufficiency also causes macular telangiectasia type 2, a retinal disease (Eade et al., 2021). Furthermore, serine is a critical component of sphingolipids. Sphingolipids, particularly gangliosides, are essential components of neuronal membranes that regulate membrane integrity, signaling, and developmental processes including neuronal maturation. Disruption of sphingolipid metabolism or synthesis can impair neuronal development and contribute to neurological disease (Schwarz et al., 1995; Olsen & Færgeman, 2017; Chinnappa et al., 2025). An increase in serine synthesis could be coordinated with an increase in sphingolipid synthesis or sphingolipid metabolism, which could have downstream effects on neurons.

Furthermore, when we tested the effect of the conditioned media from human and NHP astrocytes on neuronal maturation, we detected that NHP astrocyte conditioned media led to an increase in electrophysiological maturation and changes in the expression of neuron differentiation and maturation genes in human neurons. This result shows that species-specific rates of neuronal maturation may not be entirely cell-intrinsic but may instead be in part driven by species-specific features of astrocytes.

It is known in the literature that human neurons take longer to reach terminal maturation than NHP neurons (Marchetto et al., 2019; Somel et al., 2009; Schörnig et al., 2021; Pollen et al., 2019; Vanderhaeghen and Polleux 2023), and that human neurons eventually reach a more complex state, characterized by increased soma size, more complex branching and increased synapse numbers (Lindhout et al., 2024). We hypothesize based on the data presented here that the NHP astrocytes drive faster onset of terminal maturation of neurons, as evidenced by faster electrophysiological maturation. However, the human astrocytes release more synaptogenic and axonogenic proteins (as observed in our secretomics data), which may contribute to the increased synaptogenesis and neuronal branching reported in human neurons in other studies, suggesting different roles for human and NHP astrocytes in neurodevelopmental support. However, follow up studies at later time points to determine whether neurons exposed to human astrocyte conditioned media show an eventual increase in complexity would be necessary to conclusively determine this.

Finally, inhibition of the key step in serine synthesis led to an increase in lactate synthesis and a decrease in serine and glycine synthesis in NHP astrocytes with a more profound effect on NHP astrocytes, suggesting that NHP astrocytes are more reliant on *de novo* serine synthesis. Treating human neurons with conditioned media from NHP astrocytes treated with the inhibitor showed downregulation of neuronal differentiation genes, Together, these data suggest NHP astrocytic serine synthesis and related metabolic pathways may account for the species-specific differences in maturation effects we observed.

### Limitations of this Study

All the experiments in this paper were performed using *in vitro* models of astrocyte development. *In vitro* astrocyte features are much less developmentally mature, and the metabolic environment of these cells differs substantially from what is found *in vivo*. Although we have reanalyzed some single cell RNA-sequencing data from post-mortem human and chimpanzee astrocytes (from Jorstad et al., 2023), it is not experimentally feasible to perform metabolic studies on astrocytes from human and chimpanzee adult tissue at this time.

Furthermore, determining functional characterization of neuronal maturation is a challenge, and we assessed it with only two metrics (electrophysiology and transcriptomics). We interpreted these data as an increase in the rate of reaching terminal maturation in the NHP astrocyte conditioned media condition, but other metrics of neuronal maturation would be relevant areas of future study.

---

**Supplemental Figure 1.**
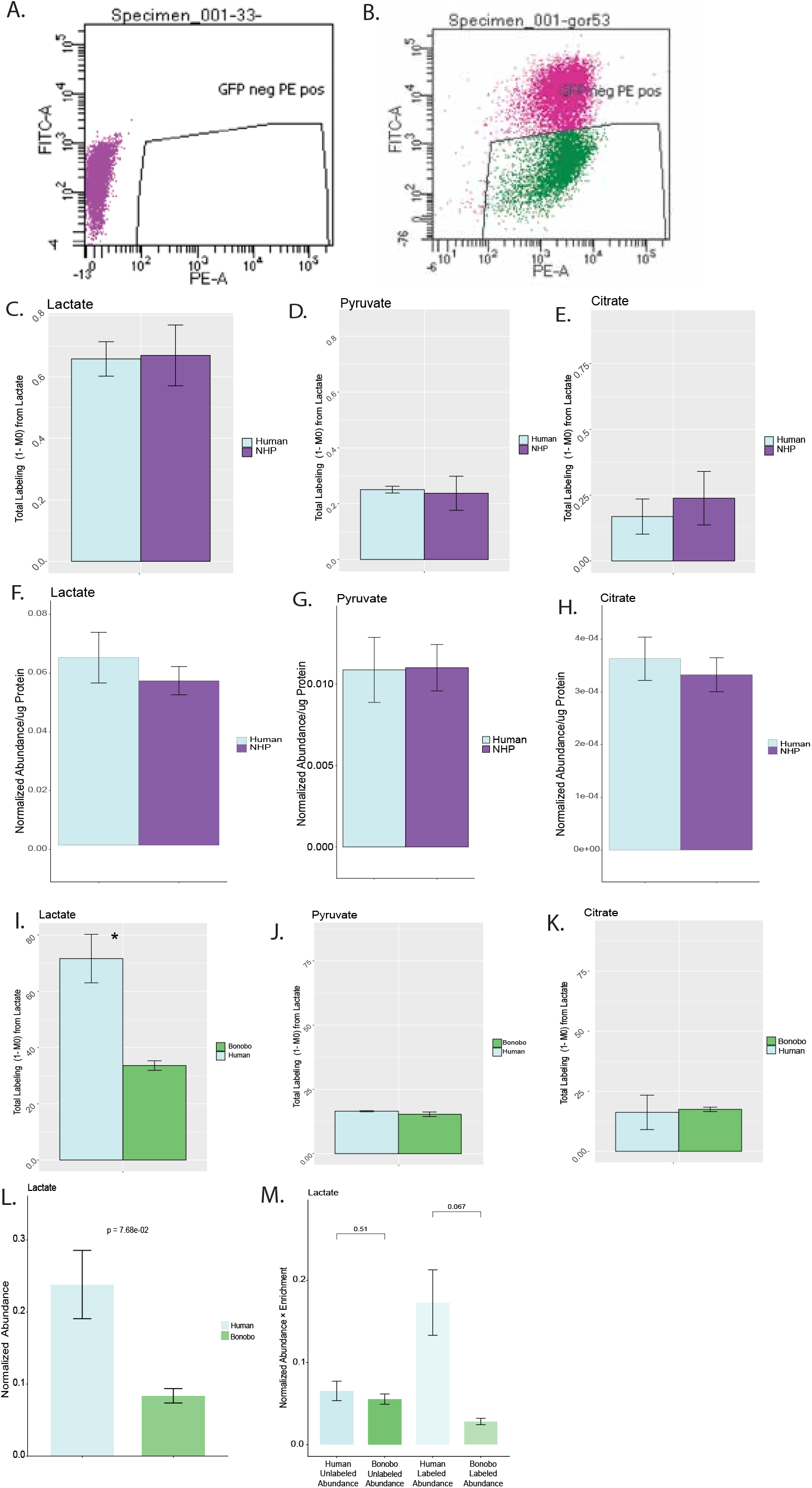
A) Example FACS gating from an unstained control human NPC line. B) Example FACS gating for PSA-NCAM + and CD44 - cells from one gorilla NPC line. C) [U-^13^C_3_] lactate tracing in one-week-old sorted neurons from three human and four NHP lines. Bars denote standard error. Technical replicates run in triplicate and averaged together per biological replicate (line). Y-axis shows 1 - M0 (or total labeling from lactate). D) Pyruvate labeling from lactate trace, as described for Supplemental Figure 1C. E) Citrate labeling from lactate trace, as described for Supplemental Figure 1C. F) Intracellular abundance of lactate normalized to norvaline (internal standard) and protein as measured by BCA from lactate trace described in Supplemental Figure 1C. G) Intracellular abundance of pyruvate, as described for Supplemental Figure 1G. H) Intracellular abundance of citrate, as described for Supplemental Figure 1G. I) [3-^13^C_1_] lactate tracing in three-week-old sorted neurons from one human and one bonobo line. Bars denote standard error. Replicates run in triplicate and represent different wells of tracing. Y-axis shows 1 - M0 (or total labeling from lactate). * = p < 0.05. J) Pyruvate labeling from lactate trace, as described for Supplemental Figure 1F. K) Citrate labeling from lactate trace, as described for Supplemental Figure 1F. L) Intracellular abundance of lactate normalized to norvaline (internal standard) and protein as measured by BCA from lactate trace described in Supplemental Figure 1I. P-value indicated. M) Abundance x enrichment values (or unlabeled and labeled abundance) for data described in Supplemental Figure 1I. T-test performed comparing human and bonobo unlabeled abundances and human versus bonobo labeled abundances.

**Supplemental Figure 2.**
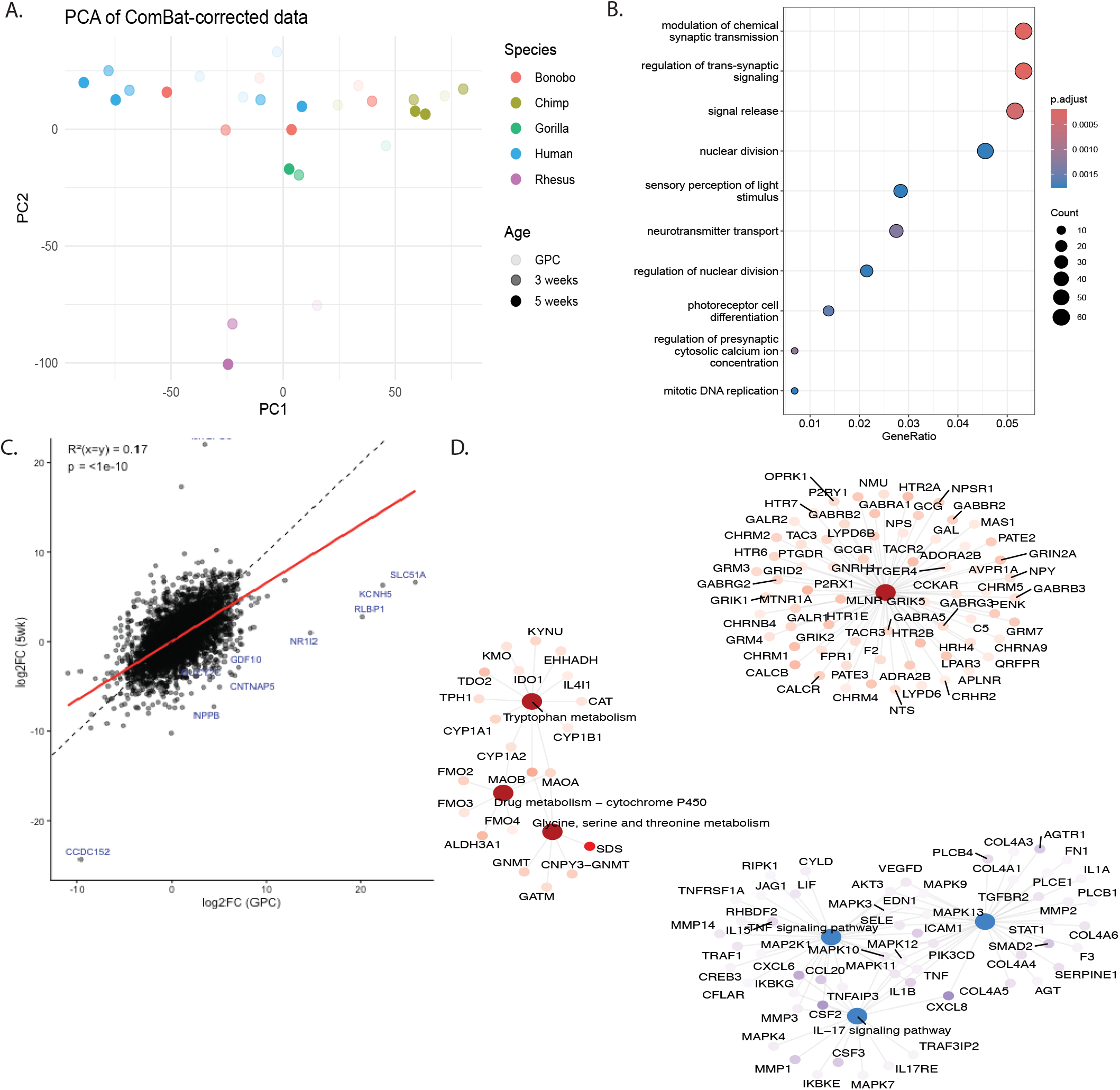
A) PCA plot of human and NHP samples as GPCs, three-week-old astrocytes and five-week-old astrocytes. Species denote color, and degree of transparency shows weeks post-differentiation. B) GO analysis for genes that are differentially expressed between humans and NHPs in both GPCs and five-week-old astrocytes. C) Plot of LFC values comparing human vs NHP in GPCs vs 5-week-old astrocytes. Dotted line shows x=y line. Red line shows the best fit line. R^2^ value and P-value show fit to the given red line. D) GSEA KEGG analysis leading edge genes within significantly enriched categories. Red gene dot indicates positive log fold change (more highly expressed in NHP five-week-old astrocytes), and blue indicates negative log fold change (more highly expressed in human five-week-old astrocytes).

**Supplemental Figure 3.**
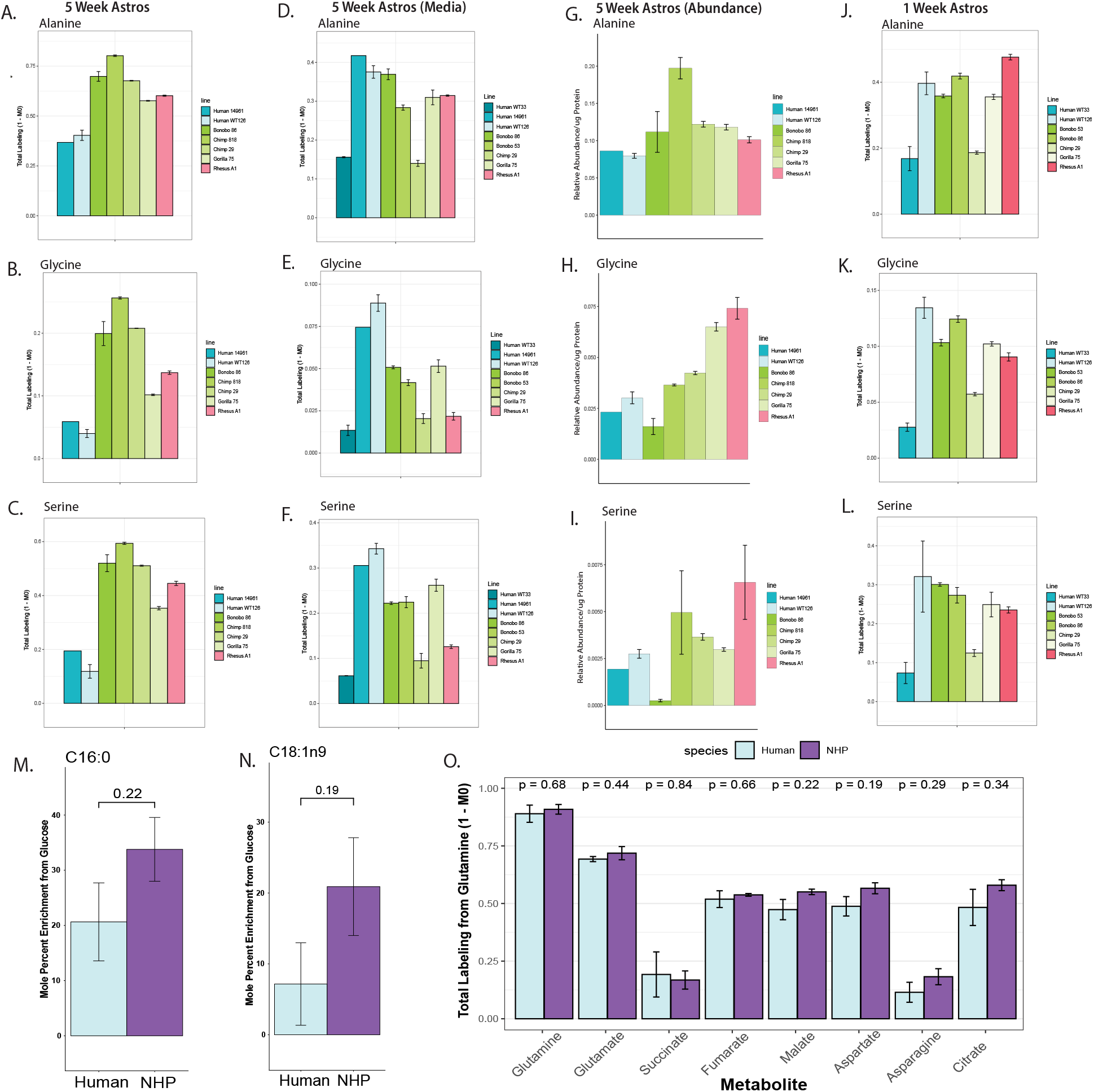
A-C) [U-^13^C_6_] glucose tracing in five-week-old astrocytes. Y axis shows 1 minus the unlabeled amount over the total amount of the given metabolite. X-axis separated by different lines, with 1 - 3 technical replicates per line (different wells of a plate). Blue shows human lines, green shows great apes and red shows rhesus macaque line (old world monkey). A shows alanine, B shows glycine and C shows serine. Error bars show standard error. Data as show in Figure 3E and F but separated by line. D-F) Labeling data as described for Supplemental Figure 5A-C but in the media from 5-week-old-astrocytes as opposed to intracellular labeling data. D shows alanine, E shows glycine, and F shows serine. G-I) Intracellular abundance data for five-week-old astrocytes. Y axis shows abundance of given metabolite normalized to an internal standard (norvaline) and protein as measured by BCA. X-axis separated by different lines, with 1 - 3 technical replicates per line (different wells of a plate). Blue shows human lines, green shows great apes and red shows rhesus macaque line (old world monkey). G shows alanine, H shows glycine and I shows serine. Bars shows standard error. J-L) Labeling data as described for A-C but for one-week old astrocytes. as opposed to intracellular labeling data. J shows alanine, K shows glycine, and L shows serine. M) Mole percent enrichment value for palmitate in human and NHP lines from [U-^13^C_6_] glucose tracing. Technical replicates are averaged together. N = 3 lines for human and N=4 lines for NHP. Bars show standard error. N) As described in N for oleic acid. O) [U-^13^C_5_] glutamine labeling data from five-week-old astrocytes. Y axis shows 1 minus the unlabeled amount over the total amount of the given metabolite. Metabolites from the TCA cycle and their branch off points are shown in order of the cycle. Technical replicates are averaged together. N = 3 for human and N=4 for NHP (biological replicates). P-values show comparison between human and NHP samples for each metabolite. Bars show standard error.

**Supplemental Figure 4.**
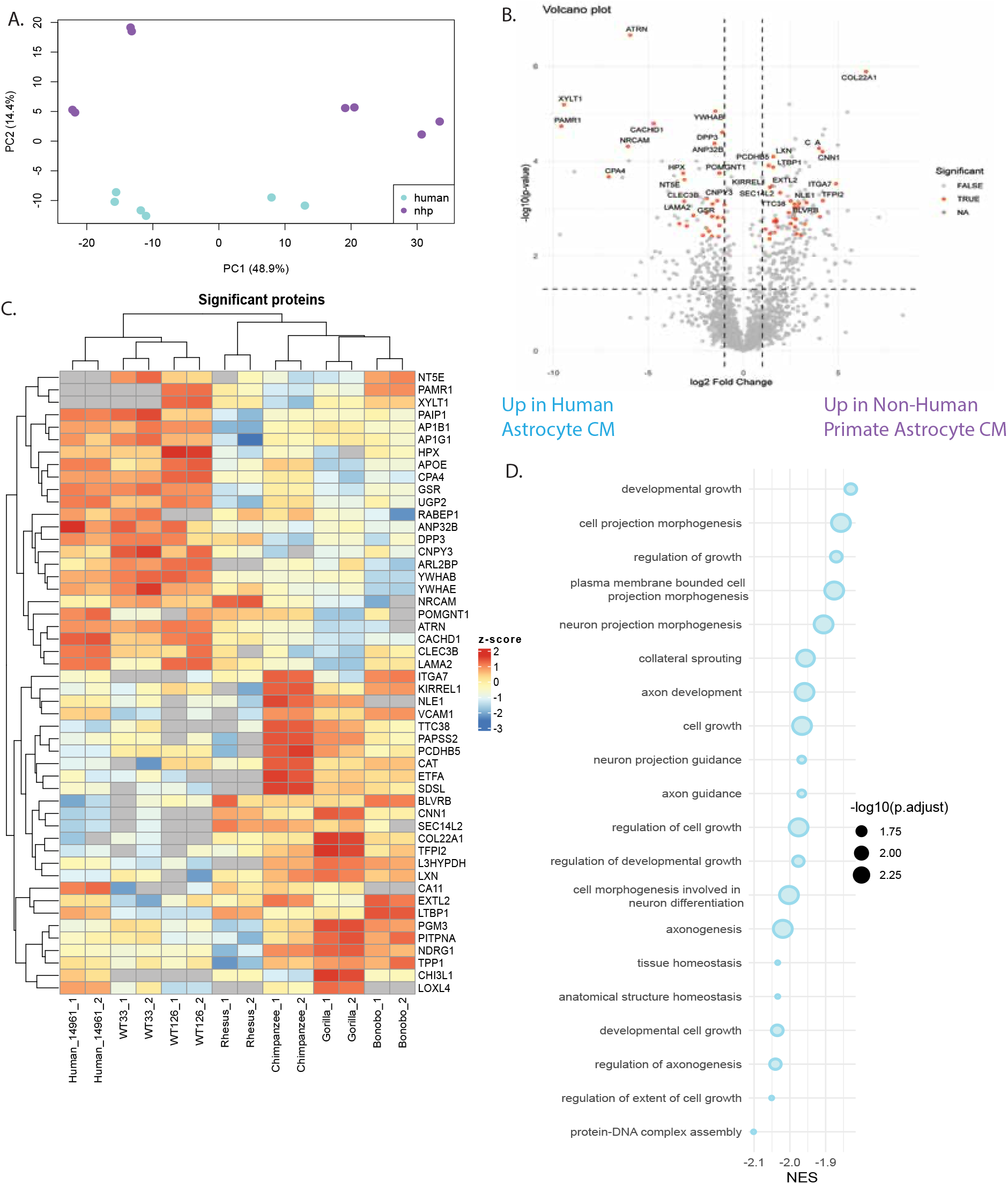
A) PCA of secretomics of human and NHP astrocytes. 3 biological replicates for human (lines) and four biological replicates for NHPs. 2 technical replicates per line. B) Volcano plot of differentially secreted proteins in humans versus NHPs. Technical replicates treated as such in MSstats in R. abs(LOG2FC) > 1 and p.adjust < 0.05 and missingPercentage < 0.3 used as cutoffs. LFC < 0 (left side) higher in abundance in human samples, LFC > 0 higher in abundance in NHP samples. C) Heatmap of top differentially secreted proteins sorted by p.adjust. Grey boxes indicate missing values. Proteins with missingPercentage > 0.3 excluded. D) GSEA for human versus NHP secretome. Top significantly differentially enriched pathways included. Pathways enriched in human secretome are colored in light blue (NES < 0).

**Supplemental Figure 5.**
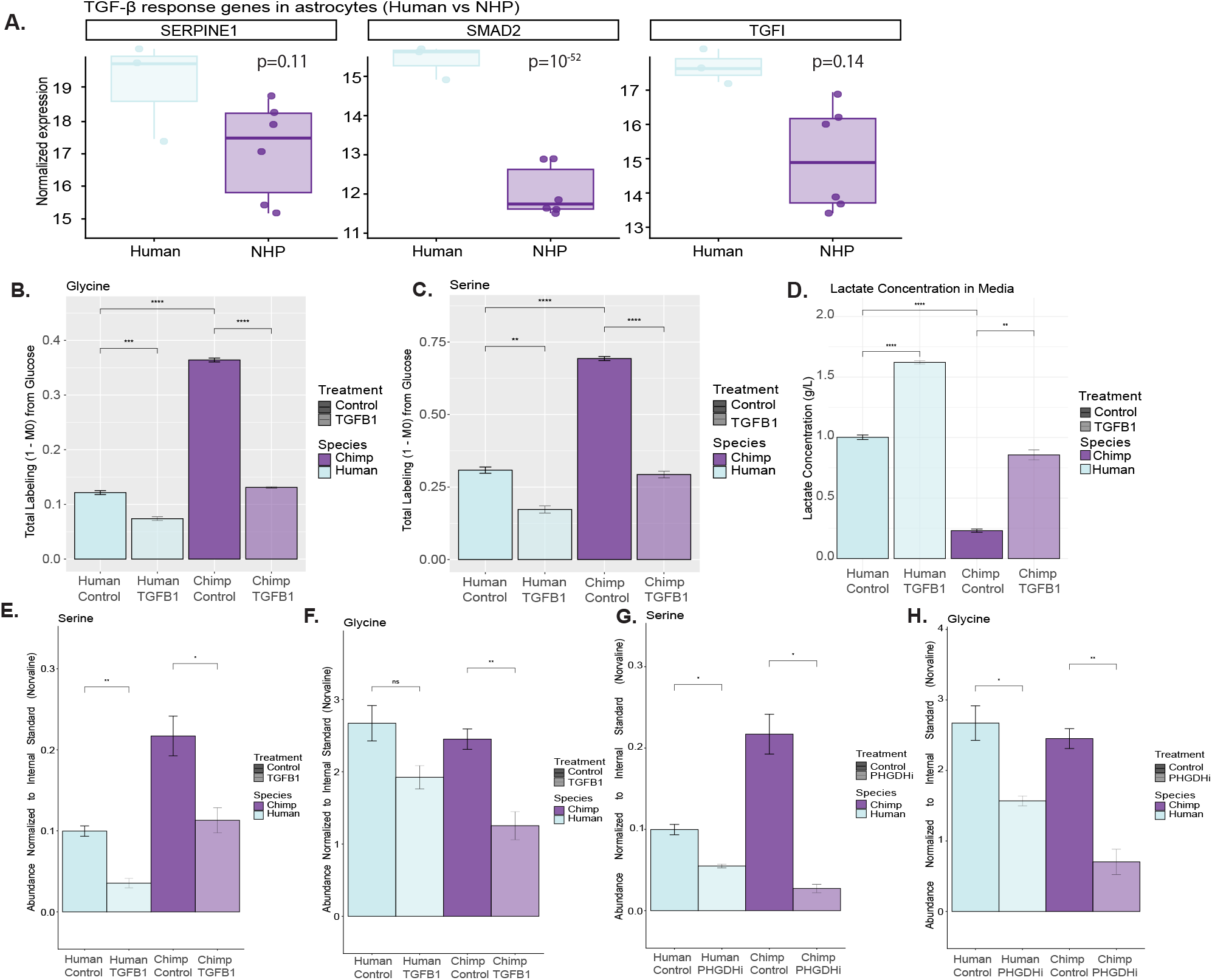
A) VST normalized and batch corrected counts for TGFβ response genes. P-values as reported from DESEQ2. B) Glycine labeling from [U-^13^C_6_] glucose. Y axis shows 1 minus the unlabeled amount over the total amount of the given metabolite. Purple indicates chimpanzee astrocyte media, and light purple shows media from chimp astrocytes treated with TGFβ1. Blue indicates human astrocyte media, and lighter blue shows media from human astrocytes treated with TGFβ1. Three wells (replicates) per condition, one line per species was used. Error bars show standard error. * = p < 0.05; ** = p < 0.01 ; *** = p < 0.001 ; **** = p < 0.0001. C) As described for Figure 5B for serine labeling. D) Lactate concentration in media as measured by YSI. Purple indicates chimpanzee astrocyte media, and light purple shows media from chimp astrocytes treated with TGFβ1. Blue indicates human astrocyte media, and lighter blue shows media from human astrocytes treated with TGFβ1. One chimpanzee line and one human line was used, and different wells were treated as replicates. Error bars show standard error. * = p < 0.05; ** = p < 0.01 ; *** = p < 0.001 ; **** = p < 0.0001. E) Intracellular abundance of serine normalized to norvaline (internal standard) from glucose trace described in Supplemental Figure 5C. Purple indicates chimpanzee astrocyte media, and light purple shows media from chimp astrocytes treated with TGFβ1. Blue indicates human astrocyte media, and lighter blue shows media from human astrocytes treated with TGFβ1. Three wells (replicates) per condition, one line per species was used. Error bars show standard error. * = p < 0.05; ** = p < 0.01 ; *** = p < 0.001 ; **** = p < 0.0001. F) As described for Supplemental Figure 5E for glycine. G) Intracellular abundance of serine normalized to norvaline (internal standard) from glucose trace described in Figure 5B. Purple indicates chimpanzee astrocyte media, and light purple shows media from chimp astrocytes treated with PHGDHi. Blue indicates human astrocyte media, and lighter blue shows media from human astrocytes treated with PHGDHi. Three wells (replicates) per condition, one line per species was used. H) As described for Supplemental Figure 5G for glycine.

## Acknowledgements

We thank all members of the laboratories of FHG, MCM and CMM for support and helpful discussions. In particular, we thank Ruth Keithley and Aurélie Laguerre for research support. We thank Dr. Pascal Gagneux and Dr. John Reynolds, members of SCS’s thesis committee, for their input and mentorship. This work was supported by the Flow Cytometry Core Facility of the Salk Institute (RRID:SCR_014839) with funding from NIH-NCI CCSG P30 CA014195. This work was also supported by the Waitt Advanced Biophotonics Core Facility of the Salk Institute (RRID:SCR_014838) with funding from NIH-NCI CCSG P30 CA014195, NIH-NIA San Diego Nathan Shock Center P30 AG068635, and the Waitt Foundation, and The Henry L. Guenther Foundation. In particular, we would like to acknowledge Elsie Quansah for performing imaging with the slide scanner and assisting with image analysis troubleshooting. This work was also supported by the Razavi Newman Integrative Genomics and Bioinformatics Core Facility of the Salk Institute (RRID:SCR_014842 and SCR_014846) with funding from NIH-NCI CCSG P30 CA014195, NIH-NIA San Diego Nathan Shock Center P30 AG068635, the NIH-NIA Liver Cancer P01 AG073084-04, the Howard and Maryam Newman Family Foundation and the Helmsley Trust. This work was supported by the Mass Spectrometry Core of the Salk Institute (RRID:SCR_014843) with funding from NIH-NCI CCSG P30 CA014195, NIH-NIA San Diego Nathan Shock Center P30 AG068635, two NIH Shared Instrumentation Grants S10-OD021815 (ThermoFisher Q-Exactive quadrupole orbitrap) and S10-OD038262 (ThermoFisher Orbitrap IQ-X Tribrid), and the Helmsley Center for Genomic Medicine. Finally, this publication includes data generated at the UC San Diego IGM Genomics Center utilizing an Illumina NovaSeq X Plus that was purchased with funding from a National Institutes of Health SIG grant (#S10 OD026929). Schematic figures were created with Biorender and some data figures were generated with GraphPad Prism. Figures were compiled and arranged in Adobe Illustrator.

This manuscript is the result of funding in part by the National Institutes of Health (NIH), award number National Institute on Aging - NIA, 5 U19 AG023122-17 (P2). It is subject to the NIH Public Access Policy. Through acceptance of this federal funding, NIH has been given a right to make this manuscript publicly available in PubMed Central upon the Official Date of Publication, as defined by NIH. The content is solely the responsibility of the authors and does not necessarily represent the official views of the NIH. Research reported in this publication was supported by the National Institute Of Aging under Award Number 5 R01 AG085634-03.

Research supported in part by the American Heart Association and the Paul G. Allen Frontiers Group Grant #19PABHI34610000/TEAM LEADER: Fred H. Gage/2019, Brinson Foundation, Grace Foundation, Freedom Together Foundation (FTF), Annette C. Merle-Smith, Robert and Mary Jane Engman Foundation, Lynn and Edward Streim, and the Ray and Dagmar Dolby Family Fund. SCS was supported by the ADRD T32 Training Grant (Grant #5T32AG066596-04) and the CARTA Fellowship.

We acknowledge support from an AHA-Allen Initiative in Brain Health and Cognitive Impairment award made jointly through the American Heart Association and The Paul G. Allen Frontiers Group (19PABH134610000) to CMM.

## Author Contributions

SCS, RS, CMM, MCM, and FHG conceived and designed the study. RS and SCS made the glial progenitor cells. KF and SCS performed astrocyte and neuron cell culture. JP, KF and SCS performed organoid cell culture. KF and SCS performed astrocyte imaging quantification. JP and SCS performed organoid slicing and staining. Imaging was done by the biophotonics core at Salk. AS and SCS did RNA extractions for PHGDH inhibition study. SCS did metabolic studies and all data analysis. RRC and CMM guided experimental design, analysis, and mass spectrometry usage. SF assisted with organoid culture and performed pilot organoid staining experiments. CMM, RS, MCM and FHG supervised studies. SCS and FHG wrote the manuscript, with input from all authors.

## Data and materials availability

All raw mass spectrometry data will be uploaded to the metabolomics workbench. Excel spreadsheets of metabolomics data are provided as spreadsheets. RNA-seq datasets and counts files will be available through GEO.

## Methods

### Cell Culture

#### Stem Cell Lines

Fibroblasts from four neurotypical humans (WT126, WT33, 14961 and MS-2C), two chimpanzees (PR00818 and PR01029), two bonobos (PR01086 and AG05253), two gorilla (PR00075 and PR00053), and one rhesus (A1) were reprogrammed to iPSCs as previously described (Linker et al., 2022, Marchetto et al., 2013, Marchetto et al., 2010, Marchetto et al., 2019, Mertens et al., 2015). Established iPS were kept in feeder-free conditions and passaged with Gentle or RelesR. Rhesus iPS were maintained in MEF conditioned media supplemented with FGF2. NPC lines were derived as previously described in Linker et al., 2022. The use of fibroblasts from chimpanzee, bonobo and gorilla were approved by the US Fish and Wildlife service, under permit MA206206. Protocols describing the use of iPSCs were previously approved by the University of California, San Diego, and the Salk Institute IRB.

#### NPC to Neuron Cell Culture

NPCs were thawed onto Matrigel or PLO/Lam plates and grown in DMEM/F12 + Glutamax + N2 + B27, with 20 ng/mL FGF2 added fresh each feed. They were split onto PLO/Lam plates and switched to neuronal differentiation media, which is as follows: DMEM/F12 + Glutamax + N2 + B27 + 20 ng/mL BDNF + 20 ng/mL GDNF + 500 ug/mL cAMP + 0.2 nM Ascorbic Acid + 1 *μ*g/mL Laminin.

#### Generation of Glial Progenitor Cells from iPS Cells

To generate glial progenitor cells, we followed the protocol as described in Santos et al. (2022) and Santos et al. (2017). In brief, we generated EBs from iPS using collagenase to generate clumps of iPS and placed them into ultra-low attachment 6 well plates. The next day, we changed the media to ScienCell Astro Media + 2% FBS and 500 ng/mL of Noggin, and 10 ng/mL of PDGF-AA. D15 after EB generation we switched the media to ScienCell Astro Media + 2% FBS + PDGF-AA at 10 ng/mL. On day 21 we dissociated the EBs using a Papain + DNase solution and plated the dissociated cells in Science Cell Astro Media + 2% FBS + 20 ng/mL FGF2 + 20 ng/mL EGF, 1 *μ*g/mL Laminin and 10 *μ*M Rock-I on Plo/Lam coated plates. The next day, cells were changed to ScienCell Astro Media + 2% FBS + 20 ng/mL FGF2 + 20 ng /mL EGF and were maintained in this media on Plo/Lam plates.

#### Generation of Astrocytes from Glial Progenitor Cells

To generate astrocytes from GPCs, we changed the media to DMEM/F12 + Glutamax + N2 + B27 – vit A + 10% FBS. On D15, we used Accutase to split cells onto uncoated plates to remove undifferentiated progenitor cells. At five weeks post-onset of differentiation, cells were considered mature.

#### IL-1β Treatment

At the five-week timepoint, astrocytes were treated with 10 ng/mL of IL-1β for five hours. They, along with a neighboring well from the same line that was not treated, were collected in Trizol for RNA-sequencing.

#### Organoid Culture

Organoid protocol is as described in Qian et al., 2018. iPS from each species were split onto MEFs. iPS + MEF cultures were fed hES media (DMEM/F12 + 20% KOSR + 1% Glutamax + 1% NEAA + 0.1% 2-Mercaptoethanol + 1% Pen Strep). Once iPS were confluent, they were split with collagenase, transferred to a 15 mL conical tube and allowed to gravity sedimentate in DMEM. They were replated in 10 cm ULA plates and fed hES media + 1:1000 rock inhibitor and 4 ng/mL of FGF2. For Days 1 – 4 EBs were fed hES + 1:5000 A83 (Tocris #2939, stock concentration 10 mM) + 1:5000 Dorsomorphin (Torcis #3093, stock concentration 10 mM). From Days 5 – 7 (or until embedding) cells were fed Forebrain Second Medium (DMEM/F12 + 1% N2 + 1% Glutamax + 1% NEAA + 1% Penn Strep + 1 uM CHIR (Torcis #4423, stock concentration 20 mM) + 1 *μ*M SB431542 (Torcis #1614, stock concentration 50 mM)).

Organoids with smooth edges were embedded in Matrigel at about the 7-day time point. Cells were fed Forebrain Second Medium from days 7 – 14.On Day 14, organoids were removed from the Matrigel using a surgical knife. Organoids were switched to forebrain third medium (DMEM/F12 + 1% N2 + 2% B27 + 1% Glutamax + 1% NEAA + 1x 2-Mercaptoethanol + 1% Pen-Strep + 2.5 *μ*g/mL Insulin (Millipore #I9278). At day 70, cells were switched to forebrain fourth medium (Neurobasal Media + 2% B27 + 1% Glutamax + 1% NEAA + 1% Pen/Strep + 0.2 mM Ascorbic Acid (Stemcell Tech #72132) + 0.5 mM cAMP (Tocris #1141) + 20 ng/mL BDNF (R&D #248-BDB) + 20 ng/ml GDNF (R&D #212-GD).

### Metabolic Tracing

#### NPC to Neuron Metabolic Tracing

NPC-derived neurons were lifted at one-week post onset of differentiation (as described above) with Accutase. Cells were stained with CD44-GFP and PSA-NCAM-PE for 45 minutes and a negative control was separated. Cells were filtered to remove clumps and FACS sorted to select for CD44-, PSA-NCAM+ cells. They were collected, spun down and replated at 200,000 cells per well of a 12 well on Plo/Lam coated plates. Cells were allowed to recover for several days and then treated with tracing media which consisted of: DMEM/F12 without glucose, 17.5 mM labeled glucose, 2 mM lactate, 3.5 mM glutamine, 1:50 B27 + VitA, 1:100 N2, 20 ng/mL BDNF (R&D #248-BDB), 20 ng/mL GDNF (R&D #212-GD), 500 *μ*g/mL cAMP (Tocris #1141), 0.2 nM Ascorbic Acid (Stemcell Tech #72132), 1 *μ*g/mL Laminin. For the 1 week and three week lactate trace, cells were fed: DMEM/F12 + 1:50 B27 + VitA, 1:100 N2, 20 ng/mL BDNF (R&D #248-BDB), 20 ng/mL GDNF (R&D #212-GD), 500 *μ*g/mL cAMP (Tocris #1141), 0.2 nM Ascorbic Acid (Stemcell Tech #72132), 1 *μ*g/mL Laminin + 3.5 mM Glutamine + 17.5 mM Glucose + 2 mM labeled lactate. After 24 hours, cells were collected as described for astrocyte tracing.

#### Astrocyte Tracing

At 5 weeks post-differentiation, astrocytes were plated at 200,000 astrocytes per well of a 12 well plate in three technical replicates per line. Two days post-plating, astrocytes were fed tracing media. Tracing media consists of DMEM/F12 without glucose, 2 mM glutamax, 1:100 N2, 1:50 B27, 1% FBS and [U-^13^C_6_] glucose for the glucose trace, and for the glutamine trace consisted of DMEM/F12, 2 mM [U-^13^C_5_] glutamine, 1:100 N2, 1:50 B27, and 1% FBS. Cells were collected as follows 48 hours following trace media addition:

First, media was collected from cells and placed into 1.5 mL Eppendorf tubes and spun down for 10 minutes at 4 degrees Celsius at max speed. Meanwhile, 1 mL 0.9% saline solution was added to each well of the twelve wells, and then removed as a rinse step. Quickly, 700 *μ*L of cold 60/40 MeOH/H2O was added to each of the twelve wells. 50 *μ*L of 100 *μ*M norvaline and 2.5 *μ*L of a fatty acid standard were added to each well. The cells were then scraped off the plate and collected in 1.5 mL Safe-Lock Eppendorf tubes. Cells were vortexed for 5 minutes. Then 75.2 *μ*L of each sample was added to a clear 96 well plate to dry out for downstream BCA protein quantification. 500 *μ*L of chloroform was added to each tube, followed by a 5-minute vortex and a 5-minute spin down at max speed at 4 degrees Celsius. 300 *μ*L of the top phase (termed the polar phase) of each Eppendorf tube was added to a mass spectrometry vial and placed open in a Centrivap at 4 degrees to dry down overnight, after which it is frozen at -80 degrees Celsius. 500 *μ*L of the bottom layer was extracted and placed into another 1.5 mL Eppendorf. This liquid was dried out under air and stored at -80 degrees Celsius.

Frozen samples were run on the mass spectrometer and analyzed using the protocol described in Green et al., 2024. Abundance values were normalized to a BCA performed on the dried down samples described above. Values for technical replicates for the same line were averaged together for downstream analyses for those plots that show humans vs non-human primates.

To analyze fatty acids, first the samples were thawed and redried under air for 10 - 15 minutes. 500 *μ*L of 2% H_2_S0_4_ in MEOH was added to each sample, and then the samples were incubated at 50 degrees Celsius for two hours. 100 *μ*L of saturated salt solution was added. 500 *μ*L MS-grade hexane was added, and samples were vortexed. After the samples settled, the top layer was placed into a new Eppendorf tube. This was then dried under air, and resuspended in 75 *μ*L of hexane, and vortexed for 10 seconds. Samples were transferred to a glass GC-MS vial with a glass insert and run on GC-MS.

#### GC-MS

Dried polar and nonpolar metabolites were processed for gas chromatography–mass spectrometry (GC-MS) as described previously by Cordes and Metallo (2021). Briefly, polar and interface metabolites were derivatized using a Gerstel MultiPurpose Sampler (MPS 2XL).

Methoxime–tert-butyldimethylchlorosilane (tBDMS) derivatives were formed by addition of 15 μL of 2% (w/v) methoxylamine hydrochloride (MP Biomedicals, Solon, OH) in pyridine and incubated at 45°C for 60 minutes. Samples were then silylated by addition of 15 μL of N-tert-butyldimethylsily-N-methyltrifluoroacetamide (MTBSTFA) with 1% tBDMS (Regis Technologies, Morton Grove, IL) and incubated at 45°C for 30 min. Derivatized polar samples were injected into a GC-MS using a DB-35MS column (30 m x 0.25 mm i.d.; Agilent J&W Scientific, Santa Clara, CA) installed in an Agilent 7890B GC system integrated with an Agilent 5977a MS. For polar metabolites, samples were injected at a GC oven temperature of 100°C which was held for 1 min before ramping to 255°C at 3.5°C/min and then to 320°C at 15°C/min and held for 3 min. For separation of the biomass metabolites the GC oven was held at 80°C for 6 min after injection, increased to 300°C at 6°C/min and held for 10 min, and finally increased to 325°C at 10°C/min and held for 4 min. Electron impact ionization was performed with the MS scanning over the range of 100 to 650 mass/charge ratio (m/z) for polar metabolites and 70-850 m/z for interface metabolites.

Nonpolar metabolites were saponified and transesterified to fatty acid methyl esters (FAMEs) by adding 500 μL of 2% H_2_SO_4_ in methanol to the dried nonpolar layer and heating at 50°C for 2 hours. One hundred uL of saturated NaCl and 500 uL of hexane were added, samples were vortexed, and the upper hexane phase was collected and transferred into a GC-MS vial. FAMEs were analyzed using a FAME Select column (100 m x 0.25 mm i.d.; Agilent J&W Scientific, Santa Clara, CA) installed in an Agilent 7890A GC system interfaced with an Agilent 5975 C MS using the following temperature program: 80C initial, increased by 20C/min to 170C, increase by 1C min/ to 204C, then 20C/min to 250C and hold for 10 min. The percent isotopologue distribution of each metabolite was determined and corrected for natural abundance using in-house algorithms that integrate the metabolite fragment ions and correct for natural isotope abundances.

#### Astrocyte Tracing with PHGDHi and TGFβ1

Cells were plated at a density of 400k cells/well of a 12 well. One day before trace media was added, PHGDH inhibitor NCT-503 (CAS 1916571-90-8) at concentration 5 uM or TGFβ1 (R&D #240-B/CF) at 1:5000 (stock concentration 50 ug/mL) was added to cells. Then 13C glucose tracing media was added to astrocytes along with fresh PHGDHi or TGFβ1 and cells were collected 48 hours later as described above and run on the mass spectrometer as described above.

### Sequencing

#### RNA seq Processing of Astrocytes and GPCs

To collect RNA, cells were washed with PBS and collected in 1 mL of Trizol at the GPC, three-week and five-week time points. To extract RNA, 200 *μ*L of chloroform were added to Trizol samples. Samples were put on ice for 5 minutes, then spun down at 12000 RCF for 15 minutes. Then the clear top layer was extracted, and an equal volume of 100% EtOH was added. RNA was extracted and purified using the RNA Clean and Concentrator Kit -5 (Zymo, Catalog # R1013). RNA was quantified using Qubit, and quality of extraction was assessed with TapeStation. Samples were submitted for sequencing at the Salk Integrative Genomics Core. Bonobo AG and Gorilla 75 GPCs and five-week astrocytes were collected and sequenced later. RNA was sequenced on a Novaseq X Plus with 20+ million reads/sample.

#### RNA seq Analysis of Astrocytes and GPCs

RNA data were aligned to the genome of the appropriate species (GCF_029289425.2, GCF_028858775.2, GCF_029281585.2,CHM13v2.0, or GCF_049350105.2). Nextflow was used for alignment, processing and QC with the following parameters: --trimmer fastp --extra_fastp_args=--trim_poly_g --pseudo_aligner kallisto -- skip_alignment true.

Data was loaded into R and non-orthologous genes or genes that had one-to-many relationships were removed, except in the case of specific genes of interest (Figure 2D). Low count and ribosomal and mitochondrial genes were removed. Raw counts were put into DESEQ2 and the formula ∼batch + nhp_human was used for GPC and 5-week analyses, and a p-value cutoff of 0.05 was used. Technical replicates were collapsed using DESEQ2 before differential expression analysis. Combat was used to perform batch correction for visualization purposes and PCA. P-values for expression (Figure 2D) were pulled from DESEQ2. GSEA was performed using KEGG with the gseKEGG function, p-value cutoff of 0.1, and minGSSize of 5. Log fold change values were derived from DESEQ2 output.

#### RNA seq Processing of CM Experiments

Three human astrocyte lines and four non-human primate astrocyte lines that were five weeks differentiated were split at the same cell density for each line. They were then transferred to neuronal media, and media was collected every 24 hours for a week from each line. Media was filtered to remove dead cells and frozen. A human NPC line (WT126) was thawed and was treated with neuronal media from one of the astrocyte lines. Neurons were collected either after one day, four days, seven days, or fourteen days in astrocyte-conditioned neuronal maturation media. A control for each time point was also collected in neuronal media without any astrocyte conditioning. RNA was extracted and purified as described for Astrocyte and GPC RNA collection. RNA was sequenced on a Novaseq X Plus with 30 - 40 M reads per sample.

#### RNA seq Analysis of CM Experiments

RNA data were aligned, processed and QC was performed using nextflow with the hg38 genome and kallisto. FASTP was used for trimming with the –trim_poly_q option. Data was loaded into R and analyzed using edgeR with the formula ∼batch + nhp_human_CM. Analysis was performed at each time point (d1, 4, 7 or 14 in conditioned media). P-value cutoff of 0.05 was used. Technical replicates were collapsed. LFC values were pulled from edgeR comparisons. GO analysis was performed using enrichGO with p-value cutoff of 0.1. For PCA plot/visualization purposes, and not differential expression analysis. Combat was used to perform batch correction for two samples that were collected at a different time.

#### RNA seq Processing of PHGDHi

One non-human primate astrocyte line (Chimp 818) that was five weeks differentiated was split at the same cell density for each line. Astrocytes were treated with a PHGDH inhibitor (NCT-503; HY-101966) for 24 hours to acclimate cells. Astrocytes were then transferred to neuronal media with a PHGDH inhibitor, and media was collected after 24 hours. Media was filtered to remove dead cells and debris. A human NPC line (WT126) was thawed, and cells were treated with neuronal media from one of the astrocyte lines. Neurons were collected after one day in astrocyte-conditioned + PHGDH inhibitor neuronal maturation media. Three controls were used; a control was collected in standard Chimp 818 astrocyte conditioned media without PHGDH inhibition, another was collected in neuronal differentiation media without astrocyte conditioning, and a third was collected from cells in neuronal differentiation media with the same concentration of the PHGDH inhibitor as was applied to the astrocytes. The third control’s media was placed in the incubator in an empty well for 24 hours to ensure similar degradation of the PHGDH inhibitor compound as was present for the astrocytes. RNA was extracted and purified as described for Astrocyte and GPC RNA collection. Different wells of NPC derived neurons were treated as replicates. Three replicates were collected for the control chimp conditioned media and PHGDHi chimp conditioned media condition and two were collected for the other two controls. RNA was sequenced using Plasmidsaurus, with 10 million reads per sample.

#### RNA seq Analysis of PHGDHi

RNA data were aligned, processed and QC was performed using nextflow with the hg38 genome and kallisto. FASTP was used for trimming with the –trim_poly_q option. Data was loaded into R and EdgeR was performed with the formula ∼treatment + conditioned_media + treatment: conditioned media. GO analysis was performed by using the function enrichGO. Genes that were significantly upregulated in the non-human primate astrocyte conditioned media versus human astrocyte conditioned media at the four day time point from the initial experiments were selected and their VST normalized counts were averaged together to create a module score for each sample. Each dot represents a replicate (different wells of NPCs). T-tests were performed in ggplot2 to compare the module scores.

### Proteomics

Five-week-old astrocytes were fed phenol-free DMEM/F12 + glutamax for 72 hours. Media was collected, frozen at -80, and given to the proteomics core for processing.

#### Proteomics Methods

Conditioned media were mixed 1:1 with a buffer containing 8 M GuHCl, 10 mM TCEP, and 40 mM CAA in 100 mM TEAB and incubated for 15 mins in the dark at 95 C to promote reduction and alkylation of disulfide bonds. The samples were then chloroform-methanol precipitated and briefly air dried. Protein precipitates were resuspended in 800 mM GuHCl in 100 mM TEAB. Proteins were digested with trypsin (1:30) overnight at 37 C. On the following day the digestion was stopped with 1% TFA, and the peptides were desalted with Empore SDB-XC StageTips (CDS Analytical). Protein amounts were normalized via a BCA assay (Pierce).

After drying in a SpeedVac, the peptides were resuspended in 0.1% FA in 5% ACN and injected into a Vanquish Neo UHPLC coupled to an Astral mass spectrometer (Thermo Fisher). Peptides were separated on a 50 cm Acclaim PepMap C18 UHPLC column (Thermo Fisher) at a flow rate of 300 nL/min. The mobile phases were 0.1% FA in water (A) and 0.1% FA in 80% ACN (B). The separation conditions were 6-12% B over 2 min, 12-28% B over 16 min, and 28-42% B over 7 min. The column was then washed at 99% B and re-equilibrated for the next sample. The mass spectrometer was operated in positive mode for data independent acquisition with a Nanospray Flex source for ionization. Full MS scans from 380-980 m/z were acquired in the Orbitrap at 240k resolution with a max injection time of 3 ms. Data independent scans were acquired in the Astral using a 2 Th isolation window, a custom AGC target of 300%, and a max injection time of 3 ms. Precursors were fragmented via HCD at 25% NCE and scans were collected from 150-2000 m/z.

LC-MS data were analyzed with CHIMERYS 4 in ProteomeDiscoverer v3.2 (Thermo Fisher) against the Uniprot Homo sapiens, Pan paniscus, Pan troglodytes, Gorilla gorilla, or Macaca mulatta reference proteome. Search settings included a mass tolerance of 10 ppm, a minimum peptide length of 7, and a maximum peptide length of 30. Full tryptic digestion was specified with a maximum of 2 missed cleavages. Modifications included fixed cysteine carbamidomethylation and variable methionine oxidation. Data was filtered at a 1% FDR at both the PSM and protein level.

Quantitation was done with the MS2 Apex strategy which uses the highest CHIMERYS coefficients from each precursor (after the 1% PSM FDR has been applied) from consecutive MS2 scans. Peptide abundances were normalized to total peptide amount, and protein abundances were calculated as the summed abundances of the connected peptides. The results were further filtered to remove proteins with only one peptide.

#### Proteomics Analysis

Proteins were summed across gene name. MS stats was run on the normalized abundance values to generate p-values comparing human and non-human primate samples. P-value cutoff of 0.05 was used. Technical replicates were entered as such in the MSstats dataProcess function. Proteins with a missingPercentage > 0.3 were not used in GSEA or the heatmap as shown in Supplemental Figure 4. GSEA GO analysis was performed on the filtered data.

### Western Blot of LDHA/LDHB

Cells were collected at the NPC state, three days post-differentiation and seven-days post differentiation from two human lines, one chimpanzee line and one gorilla line. Cell pellets were lysed with RIPA buffer and quantified using a BCA. Sample buffer was created as a master mix of BME and LDS buffer. Buffer, water and sample were mixed to a total volume of 25 *μ*uL to have an equal amount of protein between samples. Proteins were denatured at 70C for 10 minutes and then spun down. Running buffer was made by mixing 950 mL H20 with 50 mL Bolt MOPS SDS running buffer. Samples were loaded into a 4-12% Bolt BT Plus gel. Gel was run at 100 v for the first 15 minutes and then 150v until band reached the bottom of the gel. Gel was transferred using an IBlot and blocked with Odyssey blocking buffer for 1 hour at room temperature. Samples were then incubated overnight at 4 degrees Celsius in primary antibody solution. The next day, samples were washed in 0.1% PBST and shook with secondary antibody solution at room temperature for 1 hour. Blots were washed 5 times in 0.1% PBST and imaged with the Odyssey CLX.

### MEA of Human Neurons

NPCs were differentiated to neurons as described above for two weeks. They were then split with Accutase and plated onto Plo/Lam coated Maxwell 6 well plates at a density of 400k neurons/well. Neurons were allowed to recover for several days, and then a baseline recording was measured using the Maxwell. We used a firing rate threshold of 0.02 and an amplitude threshold of 20 microvolts. We recorded an activity scan for 17 minutes with 30 seconds per configuration on the full well. Then astrocyte conditioned media (collected as described above) from human and rhesus astrocytes after 24 hours of conditioning and filtering to remove cells, was added to the wells. Conditioned media was changed daily, and were recorded for several days after the baseline recording.

Wells with neurons that came off in the recording were removed, and those whose maximum % active area was less than 1.5%. Each well’s percent active area was normalized to its percent active area in the baseline recording. A t-test was performed comparing human and rhesus astrocyte conditioned media at each time point.

### Staining and Imaging of Astrocytes

To assess purity of astrocyte populations, cells were plated on 8 well slides at about 25,000 cells per well. Cells were fixed with 4% Formaldehyde solution, and then washed with PBS 3x. Cells were then treated with 0.5% Triton X-100 in PBS for one hour at room temperature. Then cells were incubated with primary antibodies in 5% horse serum in PBS overnight at 4 degrees C. The next day, cells were incubated in secondary antibody in 5% horse serum in PBS for one hour at room temperature. Cells were counterstained with DAPI for 5 minutes and then immunomount was used to mount the cells with glass slides.

For astrocytes, the following antibodies were used: rabbit anti-S100b (1:500, catalog # ab52642), mouse anti-EAAT2 (1:100, catalog # sc-365634), and chicken anti-GFAP (1:500, catalog # ab5541). The Bonobo AG line was stained at a different time than the other astrocyte lines.

Images of astrocytes were taken using an Airyscan 880 at 20x. Images were standardized across lines for laser intensity and gain to make them comparable, except for Bonobo AG because it was stained at a different time, and was brighter than the other lines, despite the same antibody concentrations.

Purity analysis for astrocytes was performed on representative 20x images. The image analysis was performed blinded to which antibody was captured and which line was being analyzed (except for Bonobo AG). Image fluorescence channels were split, and each channel was standardized to the same color balance threshold across images. With only the DAPI channel on, the images were divided into quadrants and the locations of ∼20 DAPI-stained nuclei were marked in each quadrant of the image. The exact number of DAPI nuclei marked was recorded for each quadrant. The DAPI channel was then turned off and another antibody channel was turned on. The number of marked spots with visible fluorescence was recorded. This was repeated for each channel, and the percentage of cells positive for each antibody was calculated by dividing the number of marks positive for fluorescence in that channel over the total number of marks.

### Staining and Imaging of Organoids

At each time point, five organoids were removed from the culture. They were washed in PBS 3x, fixed in PFA for 45 minutes, then washed in PO4 3x. They were then kept in 15% sucrose overnight on the shaker, then 30% sucrose overnight on the shaker. The organoids were put into molds and frozen on dry ice in TFM. They were then stored in the -80°C freezer.

Organoid blocks were then sectioned using a Cryostat at sections of 20 microns and mounted onto slides. Slides were frozen at -80°C until ready to stain. To stain, each slide border was traced with a pap-pen and then inserted into a coplin jar and washed with PBS for 5 minutes. Slides were permeabilized in a 0.1% Triton X PBS solution for 10 minutes on the shaker. 500 *μ*L of blocking buffer (5% NDS in 0.1% triton solution) were added per slide for one hour in a humidity chamber at room temperature. Primary antibody diluted in blocking buffer was added at 250 *μ*L/slide and incubated in the humidity chamber on a shaker at 4 degrees Celsius overnight. Slides were washed in PBT (0.05% Tween in PBS) 3x for five minutes each. Secondary antibody (1:200) diluted in blocking buffer was added to slides at 250 *μ*L/slide and incubated at 1 hour at room temperature in the humidity chamber. Slides were washed three times in PBT for 5 minutes/wash. Slides were then counterstained with DAPI (1:1000 in PBT) for five minutes at room temperature in the humidity chamber, and then washed 2x with PBS for five minutes per wash. Three drops of immunomount were added per slide, axnd the coverslip was mounted.

Primary antibody used were S100B (catalog # NBP2-45257, concentration 1:1000) and NFIA (catalog # 3937, active motif, concentration 1:500). Imaging was performed on an Zeiss Axioscan 7 at 20x. Imaging settings were kept consistent across all images to ensure they are comparable. In order to quantify the percent area, the pixel classifier in QuPath 6.0 was used, and several images were used as training data. To quantify the percent positive cells, we used the cell detector method in QuPath.

